# Single nucleus multi-omics links human cortical cell regulatory genome diversity to disease risk variants

**DOI:** 10.1101/2019.12.11.873398

**Authors:** Chongyuan Luo, Hanqing Liu, Fangming Xie, Ethan J. Armand, Kimberly Siletti, Trygve E. Bakken, Rongxin Fang, Wayne I. Doyle, Rebecca D. Hodge, Lijuan Hu, Bang-An Wang, Zhuzhu Zhang, Sebastian Preissl, Dong-Sung Lee, Jingtian Zhou, Sheng-Yong Niu, Rosa Castanon, Anna Bartlett, Angeline Rivkin, Xinxin Wang, Jacinta Lucero, Joseph R. Nery, David A. Davis, Deborah C. Mash, Jesse R. Dixon, Sten Linnarsson, Ed Lein, M. Margarita Behrens, Bing Ren, Eran A. Mukamel, Joseph R. Ecker

## Abstract

Single-cell technologies enable measure of unique cellular signatures, but are typically limited to a single modality. Computational approaches allow integration of diverse single-cell datasets, but their efficacy is difficult to validate in the absence of authentic multi-omic measurements. To comprehensively assess the molecular phenotypes of single cells in tissues, we devised single-nucleus methylCytosine, Chromatin accessibility and Transcriptome sequencing (snmC2T-seq) and applied it to post-mortem human frontal cortex tissue. We developed a computational framework to validate fine-grained cell types using multi-modal information and assessed the effectiveness of computational integration methods. Correlation analysis in individual cells revealed distinct relations between methylation and gene expression. Our integrative approach enabled joint analyses of the methylome, transcriptome, chromatin accessibility and conformation for 63 human cortical cell types. We reconstructed regulatory lineages for cortical cell populations and found specific enrichment of genetic risk for neuropsychiatric traits, enabling prediction of cell types with causal roles in disease.

## INTRODUCTION

Single-cell transcriptome, cytosine DNA methylation (mC) and chromatin profiling techniques have been successfully applied for cell-type classification and studies of gene expression and regulatory diversity in complex tissues (Ecker et al., 2017; Kelsey et al., 2017). The broad range of targeted molecular signatures, as well as technical differences between measurement platforms, presents a challenge for integrative analysis. For example, mouse cortical neurons have been studied using single-cell assays that profile RNA, mC or chromatin accessibility (Luo et al., 2017; Preissl et al., 2018; Tasic et al., 2016, 2018; Zeisel et al., 2015), with each study reporting its own classification of cell types. Although it is possible to correlate the major cortical cell types identified by transcriptomic and epigenomic approaches, it remains unclear whether fine subtypes can be effectively integrated across different datasets and between modalities. Recently, computational methods based on Canonical Correlation Analysis (CCA, e.g. Seurat3) (Stuart et al., 2019), mutual nearest neighbors (MMN, e.g. Scanorama) (Hie et al., 2019) or matrix factorization (e.g. LIGER) (Welch et al., 2019) have been developed to integrate molecular data types. However, validating the results of computational integration requires multi-omic reference data comprising different types of molecular measurements made in the same cell.

Single-cell multi-omic profiling provides a unique opportunity to evaluate cell types classification using multiple molecular signatures (Kelsey et al., 2017). Most single-cell studies rely on clustering analysis to identify cell types. However, it is challenging to objectively determine whether the criteria used to distinguish cell clusters are statistically appropriate and whether the resulting clusters reflect biologically distinct cell types (Mukamel and Ngai, 2019). We reasoned that genuine cell types should be distinguished by concordant molecular signatures of cell regulation at multiple levels, including RNA, mC and open chromatin, in individual cells. Moreover, multi-omic data can uncover subtle interactions among transcriptomic and epigenomic levels of cellular regulation.

Existing methods for joint profiling of transcriptome and mC, such as scM&T-seq and scMT-seq, rely on physical separation of RNA and DNA followed by parallel sequencing library preparation (Angermueller et al., 2016; Clark et al., 2018; Hu et al., 2016). Generating separate transcriptome and mC sequencing libraries leads to a complex workflow and increases cost. Moreover, it is unclear if these methods can be applied to single nuclei, which contain much less polyadenylated RNA than whole cells. Since the cell membrane is ruptured in frozen tissues, the ability to produce robust transcriptome profiles from single nuclei is critical for applying a multi-omic assay for cell-type classification in frozen human tissue specimens.

Here we describe two single nucleus multi-omic methods that do not require physical separation of RNA and DNA. Single-nucleus methylCytosine & Transcriptome sequencing (snmCT-seq) captures mC and transcriptome profiles from single cells/nuclei, whereas snmC2T-seq (single-nucleus methylCytosine, Chromatin accessibility and Transcriptome sequencing) simultaneously interrogates transcriptome, mC and chromatin accessibility, based on NOMe-seq (Clark et al., 2018; Guo et al., 2017; Kelly et al., 2012; Pott, 2017). We applied both methods to cultured human cells and postmortem human frontal cortex tissues. We further generated an additional 23,005 single-nucleus, droplet-based RNA-seq profiles and 12,557 single-nucleus, snATAC-seq-based open chromatin profiles using frozen human frontal cortex tissue (Preissl et al., 2018). Using this comprehensive multimodal dataset, we developed computational strategies to tackle two challenges in single-cell biology: 1) how to assess the statistical and biological validity of clustering analyses, and 2) how to validate computational approaches to integrate multiple single-cell data types. We then performed integrated analyses of single-cell methylomes for the human frontal cortex comprised of 15,030 cells, including two multi-omic data sets generated by snmC2T-seq and the previously published sn-m3C-seq, a method to simultaneously profile chromatin conformation and mC (Lee et al., 2019). These large datasets enabled the identification of gene regulatory diversity for 63 finely defined brain cell types at an unprecedented level of integration using four levels of molecular signatures (i.e. transcriptome, methylome, chromatin accessibility, and conformation) to define their unique regulatory genomes with cell-type specificity and link them to genetic disease risk variants.

## RESULTS

### Joint analysis of RNA and DNA methylome with molecular partitioning

Simultaneous DNA methylcytosine and transcriptome sequencing using mCT-seq allows RNA and DNA molecules to be molecularly partitioned by incorporating 5’-methyl-dCTP instead of dCTP during reverse transcription of RNA (Figure 1A). We treated single nuclei with Smart-seq or Smart-seq2 reactions for *in situ* cDNA synthesis and amplification of full-length cDNA (Picelli et al., 2013; Ramsköld et al., 2012). Replacing dCTP by 5’-methyl-dCTP results in fully cytosine-methylated double-stranded cDNA amplicons. Following bisulfite treatment converting unmethylated cytosine to uracil, sequencing libraries containing both cDNA- and genomic DNA-derived molecules were generated using snmC-seq2 (Luo et al., 2017, 2018). With this strategy, all sequencing reads initially derived from RNA are completely cytosine methylated and do not show C to U sequence changes during bisulfite conversion. By contrast, more than 95% of cytosines in mammalian genomic DNA are unmethylated and converted by sodium bisulfite to uracils that are read during sequencing as thymine (Lister et al., 2009). In this way, sequencing reads originating from RNA and genomic DNA can be distinguished by their total mC density. Since 70-80% of CpG dinucleotides are methylated in mammalian genomes, we used the read-level non-CG methylation (mCH) to uniquely partition sequencing reads into RNA or DNA bins. Specifically, we expect the level of mCH for all RNA-derived reads to be greater than 90%, while for DNA derived reads the level is no more than 50% even considering the enrichment of mCH in adult neurons (Lister et al., 2013). Using this threshold, only 0.04% of single-cell methylome reads were misclassified as transcriptome reads and only 0.23% ± 0.03% of single-cell RNA-seq reads were misclassified as methylome reads. These results show that RNA- and DNA-derived mCT-seq reads can be effectively separated.

**Figure 1.**
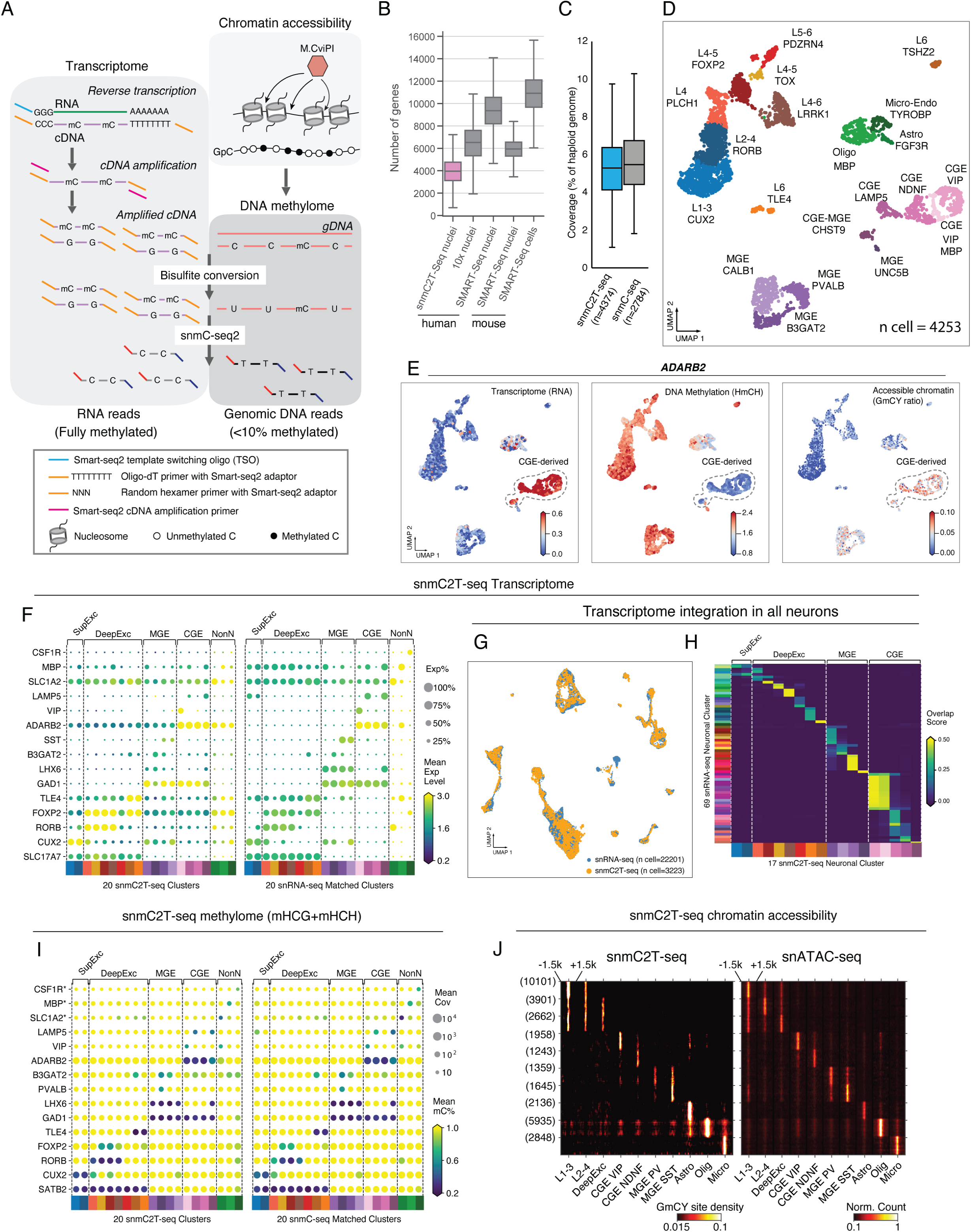
snmC2T-seq generates single-nucleus multi-omic profiles of the human brain. (**A**) Schematic diagram of snmC2T-seq. (**B**) Boxplot comparing the number of genes detected in each cell/nucleus by different single-cell or single-nucleus RNA-seq technologies. (**C**) Boxplot comparing the genome coverage of single-nucleus methylome between snmC2T-seq and snmC-seq. (**D**) UMAP embedding of human frontal cortex snmC2T-seq profiles. (**E**) UMAP embedding of transcriptome, methylome and chromatin accessibility profiled by snmC2T-seq for *ADARB2*. From left to right, the cells are colored by gene expression (CPM, counts per million), non-CG DNA methylation (HmCH ratio normalized per cell) and chromatin accessibility (MAGIC imputed GmCY ratio, see methods). (**F**) Comparison of marker gene expression between clusters identified using snmC2T-seq and matching clusters identified using snRNA-seq. The matching clusters were merged from original snRNA-seq clusters based on cell integration and label transfer (see methods). Dot sizes represent the fraction of cells with detected gene expression. Dot colors represent the mean expression level across the cells with detected gene expression. (**G**) UMAP embedding of snmC2T-seq transcriptome and snRNA-seq cells after integration. (**H**) Confusion matrix comparing snmC2T-seq clusters to snRNA-seq clusters. The plot is colored by overlapping scores between clusters. (**I**) Comparison of marker gene non-CG methylation (HmCH) between clusters identified using snmC2T-seq and matching clusters identified using snmC-seq. Dot sizes represent the mean cytosine coverage per cell. Dot colors represent the mean HmCH ratio. *For non-neuronal cell markers, gene body CG methylation (HmCG) levels were compared between snmC2T-seq and snmC-seq. (**J**) Comparison of chromatin accessibility profiled by snmC2T-seq and snATAC-seq at cell-type-specific open chromatin sites. The left and right heatmaps show the density of methylated GCY sites and the density of ATAC-seq reads, respectively.

To test the efficacy of these methods, we first applied mCT-seq to either single whole cells (scmCT-seq) or single nuclei (snmCT-seq) of cultured human H1 embryonic stem cells and HEK293 cells (Table S1-S2). scmCT-seq transcriptome profiling detected 4,220 ± 1,251 genes from single whole cells using exonic reads while snmCT-seq detected 4,531 ± 1,888 genes using both exonic and intronic reads (Figure S1A). Similar to previously reported single-nuclei RNA-seq datasets, a minor fraction (17.3 ± 6.1%) of snmCT-seq transcriptome reads were mapped to exons, whereas 68.1 ± 15.2% of scmCT-seq reads were mapped to exons (Figure S1B). Transcriptome reads accounted for 22.2 ± 13.6% and 9.2 ± 6.5% of all mapped reads for scmCT-seq and snmCT-seq, respectively (Figure S1C). The two human cell types were clearly separated by their transcriptomic signatures measured using either scmCT-seq or snmCT-seq (Maaten and Hinton, 2008) (Figure S1D-E). Further, scmCT-seq or snmCT-seq profiles recapitulate H1 or HEK293 specific gene expression signatures (Figure S1F).

To assess whether the two cell types could be distinguished using mC signatures derived from snmCT-seq, tSNE was performed using the average CG methylation (mCG) level of 100 kb non-overlapping genomic bins measured from single cells or nuclei (Figure S1G-H). As exemplified by the NANOG and CRNDE loci (Figure S1I), both single-cell multi-omic assays produced mC profiles highly consistent with data generated from bulk methylomes (Ellis et al., 2012). scmCT-seq and snmCT-seq identified global mC differences between H1 and HEK293T cells, showing that H1 cells are more methylated in both CG (83.6%) and non-CG (1.3%) contexts compared with HEK293T cells (mCG: 60.1%, no significant mCH detected, Figure S1J-M) (Lister et al., 2009). To examine whether local mC signatures can be recapitulated in scmCT-seq and snmCT-seq profiles, we identified differentially methylated regions (DMRs) from bulk H1 and HEK293 methylomes. Plotting mCG levels measured using scmCT-seq and snmCT-seq across DMRs showed highly consistent patterns compared to bulk cell methylomes (Figure S1N-O).

### Multi-omic profiling of postmortem human brain tissue with snmC2T-seq

Next we extended snmCT-seq to include a measure of chromatin accessibility by incorporating the Nucleosome Occupancy and Methylome-sequencing assay (NOMe-seq, Figure 1A) (Clark et al., 2018; Guo et al., 2017; Kelly et al., 2012; Pott, 2017). In the snmC2T-seq assay, regions of accessible chromatin are marked by treating bulk nuclei with the GpC methyltransferase M.CviPI prior to fluorescence-activated sorting of single nuclei into the reverse transcription reaction (Figure 1A). We generated snmC2T-seq profiles from 4,358 single nuclei isolated from postmortem human frontal cortex tissue from two young male donors (21 and 29 years old, Table S3-4). The data quality was similar to datasets generated from cultured human cells with respect to the fraction of sequencing reads mapped to the transcriptome (Figure S2A), the fraction of transcriptome reads mapped to introns and exons (Figure S2B) and the number of genes detected (Figure 1B and Figure S2C). Compared with snmC-seq and snmC-seq2 data generated from human single nuclei (Luo et al., 2017, 2018), the DNA methylome component of snmC2T-seq had comparable genomic coverage (Figure 1C), mapping efficiency (Figure S2D), and showed only moderately reduced library complexity (Figure S2E) with similar coverage uniformity (Figure S2F-G).

To compare each data modality profiled by snmC2T-seq with their corresponding single modality assays, we first identified 20 cell types by jointly using transcriptome, methylome and chromatin accessibility. We used RNA abundance across gene body for the transcriptome, mCH and mCG level of chromosome non-overlapping 100kb-bins, and binarized NOMe-seq signal of 5kb bins for chromatin accessibility (See methods). In each modality, we identified highly variable features separately and calculated Principle Components, we then concatenated the PCs together as the input features for joint clustering and UMAP (McInnes et al., 2018) visualization of the three data types (Figure 1D-E). These cell types were effectively separated by performing dimensionality reduction using each individual data type (Figure S2H-J). The comparison of homologous clusters between snmC2T-seq transcriptome and snRNA-seq (Table S5) shows a robust global correlation - Pearson r = 0.82 for both PV-expressing inhibitory neurons (MGE_PVALB, p = 1×10^-145^) and superficial layer excitatory neurons (L1-3 CUX2, p = 3×10^-301^) (Figure S2K-L). Moreover, highly consistent expression patterns of cell-type signature genes were observed (Figure 1F).

To test whether snmC2T-seq transcriptome data can be integrated with snRNA-seq, we performed joint embedding and clustering of snRNA-seq and the transcriptome component of snmC2T-seq (Figure 1G). The joint clustering confirmed that the cell types identified using the snmC2T-seq transcriptome are strongly correlated with the cell types found using snRNA-seq (Figure 1H). Similar to the transcriptome, both mCH and mCG profiles correlate strongly between methylomes generated with snmC2T-seq and snmC-seq2 either globally (Figure S2M-N) or at cell type-specific signature genes (Figure 1I).

The presence of high levels of mCH in the human brain confounds the analysis of chromatin accessibility using methylation at GpC sites (GmC). However, we found that in GCT and GCC sequence contexts, GmC introduced by M.CviPI greatly surpasses the levels of native methylation by 6.4 and 16-fold, respectively (Figure S2O). Thus for snmC2T-seq, we focused our analyses of chromatin accessibility on GmC at GCY (Y=C or T) sites in the genome. We further developed a computational strategy to effectively identify open chromatin regions using the frequency of significantly methylated GCY (GmCY) sites. Chromatin accessibility measured by the frequency of GmCY sites correlates closely with snATAC-seq signal at cell-type specific open chromatin sites (Figure 1J and Figure S2P-Q, p-value < 2.2 x 10^-308^). In addition, open chromatin regions identified with GmCY frequency overlapped substantially with regions found using snATAC-seq (Fig.S2R-S). In summary, snmC2T-seq can simultaneously profile transcriptome, methylome and chromatin accessibility in single nuclei, accurately recapitulating cell-type signatures for each data type.

### Multi-omic integration of chromatin conformation, transcriptome, methylome and chromatin accessibility

We then generated snmC2T-seq and snmCT-seq profiles for human frontal cortex from two independent donors. These data were combined with previously published human frontal cortex datasets (Table S3): sn-m3C-seq which simultaneously profile mC and chromatin conformation (Lee et al., 2019) and snmC-seq methylomes for 2,784 single neurons (Luo et al., 2017). These datasets can be readily integrated by joint embedding and clustering using single-nucleus methylomes as the common framework (Figure 2A). To identify both major cell types and subtypes of frontal cortex, we performed joint clustering of 15,030 single-cell methylomes generated by snmC-seq (n=5,131), snmC-seq2 (n=1,304), snmC2T-seq (n=4,358) and sn-m3C-seq (n=4,238) (Table S6). We used an iterative clustering approach to identify 20 major cell populations including 9 excitatory neuron types, 8 inhibitory neuron types and 3 non-neuronal cell types in the first round of clustering (Figure 2B-C). A second round of iterative clustering of each major cell types identified 63 cell subtypes, including 19 excitatory neuronal subtypes, 33 inhibitory neuronal subtypes and 11 non-neuronal cell subtypes (Figure 2B-C). Each fine-grained cell subtypes can be distinguished from any other cell type by at least 10 mCH signatures genes for neuronal clusters, or 10 mCG signatures genes for non-neuronal clusters. Consistent with our previous results (Luo et al., 2017) as well as transcriptomic studies (Hodge et al., 2019), we found greater diversity among human cortical inhibitory neurons than excitatory cells (Figure 2C). The methylome data generated by these diverse multi-omic methods and from multiple donors were uniformly represented in major cell type and subtype clusters (Figure 2D).

**Figure 2.**
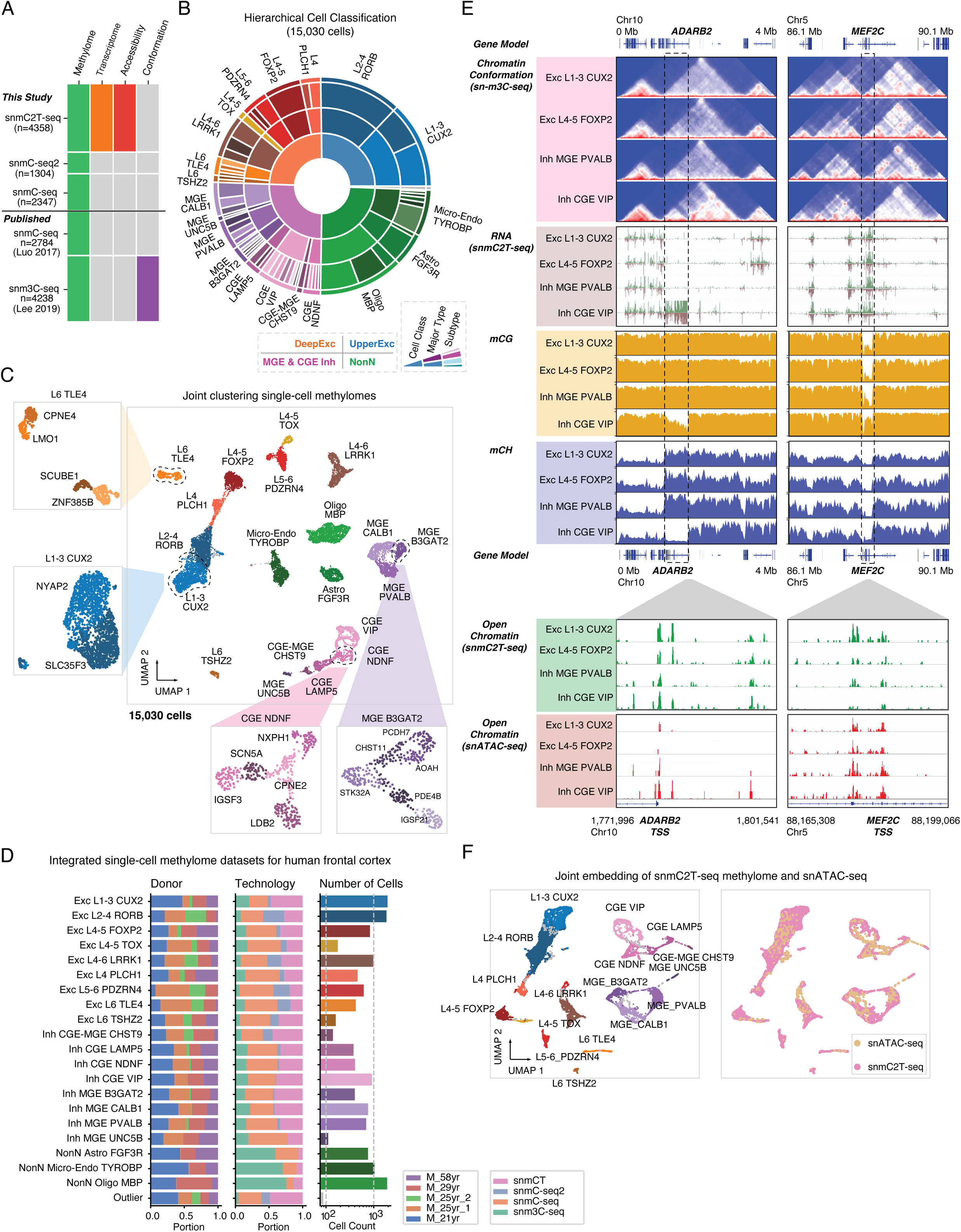
Integrated epigenomic atlas of the human frontal cortex. (**A**) Methylome based technologies and datasets included in the integrative analysis. (**B**) Sunburst visualization of the two-level methylome ensemble clustering analysis. The 4 cell classes (inmost ring) and 20 major cell types (middle ring and outer annotation) are identified in level 1 analysis, the 63 subtypes are identified in level 2 analysis. (**C**) UMAP embedding of 15,030 cells colored and labeled by major cell types from level 1 analysis. Several examples of level 2 analysis are shown in insets with UMAP colored and labeled by subtypes. (**D**) Donor (left) and technology (middle) composition and cell count (right) of each major cell type. (**E**) Browser views of multi-modal data integration for *ADARB2* and *MEF2C* gene in four major cell types. (**F**) UMAP embedding of the cross-modality integration of snmC2T-seq methylome and snATAC-seq profiles. The left panel is colored and labeled by level 1 major cell types; the right panel is color and labeled by the technologies.

We next performed integration of single-cell methylome and snATAC-seq (Table S7) profiles by transferring the cluster labels defined by mC into ATAC-seq cells using a nearest neighbor approach (Haghverdi et al., 2018) that was adapted for epigenomic data and implemented in a new software package (https://github.com/mukamel-lab/SingleCellFusion, see methods). For each cell population, we reconstructed four types of molecular profiles: transcriptome (from snmC2T-seq), methylome (from snmC-seq1/2, snmCT-seq and sn-m3C-seq), chromatin accessibility (from snmC2T-seq mGCY frequency or snATAC-seq) and chromatin conformation (from sn-m3C-seq) (Figure 2E). This integrative analysis revealed extensive correlations across epigenomic marks at cell-type signature genes. For example, *ADARB2* is a signature gene of inhibitory neurons derived from the caudal ganglionic eminence (CGE). In CGE-derived VIP neurons, *ADARB2* was associated with abundant transcripts, reduced mCG and mCH, and distinct chromatin interactions compared with other neuron types (Figure 2E). In contrast, in VIP neurons the *MEF2C* locus showed lower transcript abundance (TPM - L1-3 CUX2: 75.8, L4-5 FOXP2: 80.2, MGE PVALB: 77.5, CGE VIP: 49.0), reduced chromatin interaction, and more abundant gene body mCG (Figure 2E). Although nearly identical open chromatin sites were identified at the promoter regions of *ADARB2* and *MEF2C* using GmCY frequency and snATAC-seq, the two methods revealed distinct cell-type specificity of chromatin accessibility. At the *ADARB2* promoter, snATAC-seq but not the GmCY frequency profile showed enriched chromatin accessibility in VIP neurons. However, at *MEF2C* promoter, the GmCY frequency indicated a depletion of open chromatin in VIP neurons which is more consistent with the reduced gene expression and increased gene body mCG in this inhibitory cell population. The cause of these differences in measures of chromatin accessibility is not clear, and further work is needed to clarify their respective sensitivity and biases.

### snmC2T-seq identifies RNA and mC signatures of neuronal subtypes

Joint clustering of 15,030 single-cell methylomes allowed determination of fine-grained brain cell subtypes with a sensitivity comparable to snRNA-seq (Figure 2B-C). For example, we identified 15 subtypes of CGE-derived inhibitory neuron using single-cell methylomes, whereas 26 subtypes were identified by snRNA-seq (Hodge et al., 2019). To ask whether snmC2T-seq can recapitulate the molecular signatures of neuronal subtypes, we first integrated snmC2T-seq transcriptome with snRNA-seq datasets for inhibitory neurons using Scanorama followed by joint clustering (Figure 3A-B). Individual nuclei profiled with snmC2T-seq transcriptome and snRNA-seq were uniformly distributed across joint clusters corresponding to inhibitory neuron subpopulations (Figure 3B-C), suggesting that the snmC2T-seq transcriptome recapitulates the full range of inhibitory neuron diversity. Similarly, integration of snmC2T-seq transcriptomes and snRNA-seq for excitatory neurons and non-neuronal cells showed that brain cell type diversity across all cell classes can be recapitulated from the snmC2T-seq transcriptome profiles. (Figure S3D-I). We further compared the expression of a panel of signature genes for inhibitory neuron subpopulations and found that snmC2T-seq transcriptome and snRNA-seq identified highly consistent expression patterns (Figure 3D). Lastly, we identified cell-type marker genes across inhibitory neuronal populations using transcriptome profiles generated with either snmC2T-seq or snRNA-seq (Table S8). Analysis of the marker genes using a database curated for neuronal functions - SynGO (Koopmans et al., 2019) revealed consistent enrichment in ontological categories associated with synaptic signaling and synapse organization for inhibitory neuron marker genes identified with both snmC2T-seq transcriptome and snRNA-seq data (Figure 3E).

**Figure 3.**
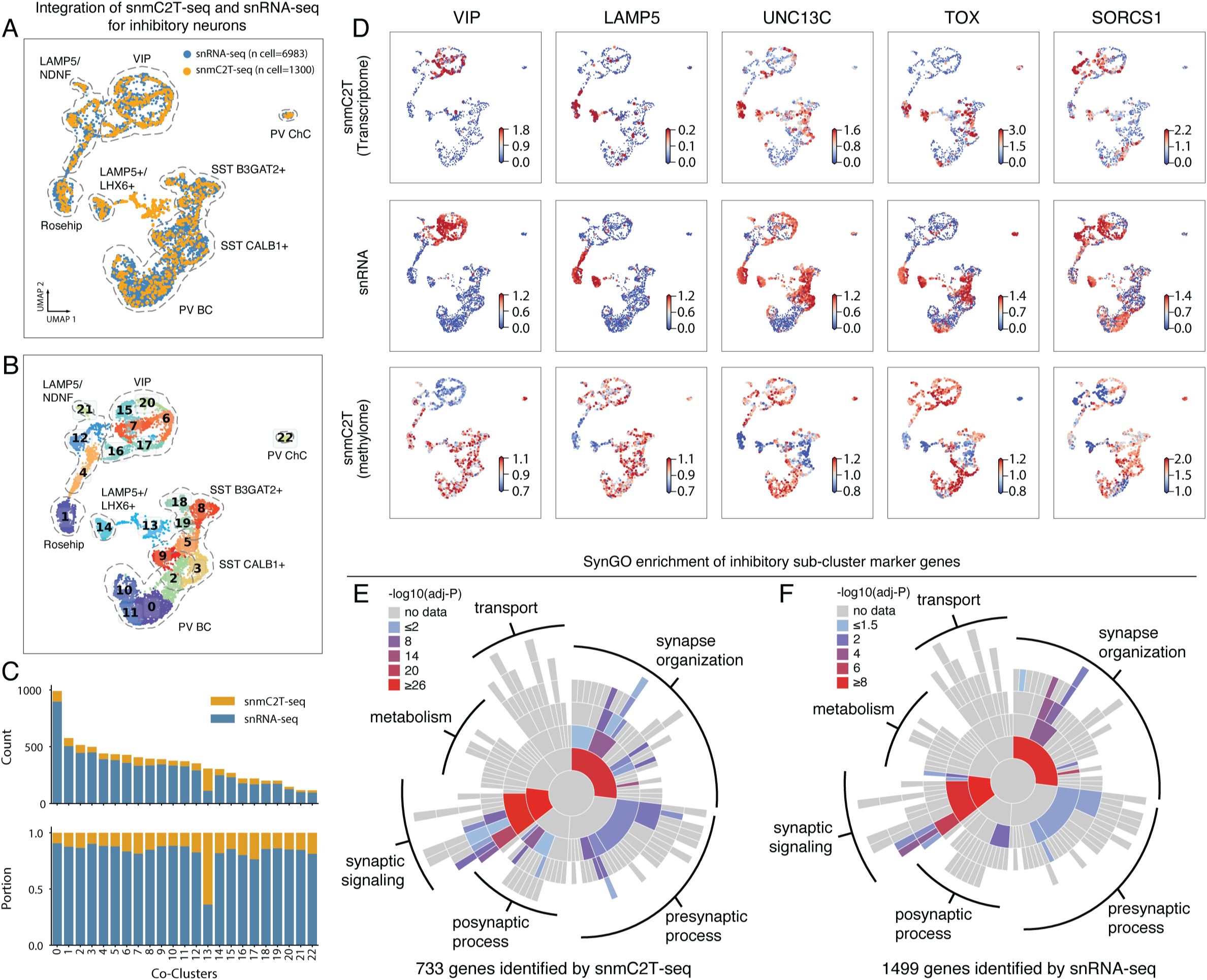
snmC2T-seq identifies RNA and mC signatures of neuronal subtypes. (**A-B**) UMAP embedding of snmC2T-seq transcriptome and snRNA-seq for all the inhibitory neurons after MNN-based integration with the cells colored by technology (A) and joint clusters (B). (**C**) The composition of cells profiled by snmC2T-seq and snATAC-seq in inhibitory neurons joint clusters (same cluster IDs as shown in (B)). The upper and lower bar plots show the counts and portion of cells profiled by the two technologies in each joint cluster, respectively. (**D**) Normalized expression and gene body mCH rate of inhibitory neuron subtype marker genes quantified using snmC2T-seq transcriptome and snRNA-seq. (**E-F**) Sunburst visualization of inhibitory cell type marker genes enrichment in SynGO biological process terms. Each sector is a SynGO term colored by −log10(adjusted P-value) of snmC2T-seq transcriptome marker genes (E) or snRNA-seq marker genes (F) enrichment.

### Paired RNA and mC profiling enables cross-validation and quantification of over-/under-splitting for single-cell clusters

A fundamental challenge for single-cell genomics is to objectively determine the number of biologically meaningful clusters in a dataset (Mukamel and Ngai, 2019). Cross-dataset integration can be used to assess cluster robustness, but it may be limited by systematic differences between the datasets or modalities used (Crow et al., 2018). To address this, we devised a novel cross-validation procedure using matched transcriptome and DNA methylation information to estimate the number of reliable clusters supported by both modalities in snmC2T-Seq data (3,898 neurons, Figure 4A). The cells were first clustered with different resolutions using mC information followed by testing how well each clustering is supported by the matched transcriptome profiles, as represented by the mean squared error (MSE) between the RNA expression profile of individual cells and that of cluster centroids as a function of the number of clusters (Figure 4B-C). Increased cluster number monotonically reduces MSE in the training set, whereas overclustering (more than 20 clusters) leads to an increase in MSE for the test set (Figure 4B). Using this approach, we found a range of 20-50 clusters with strong multimodal support in the current snmC2T-seq dataset (Figure 4B,C).

**Figure 4.**
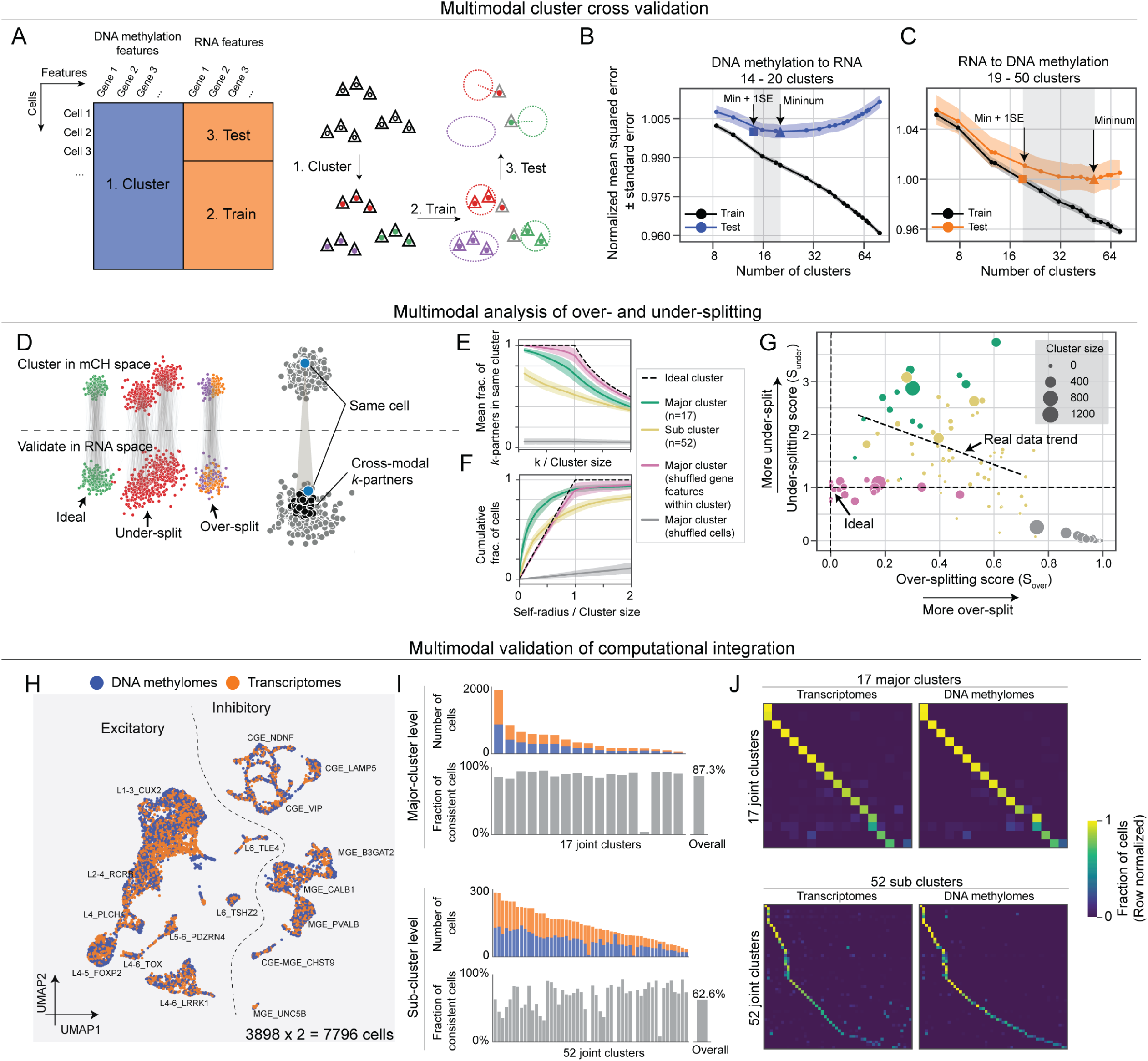
Integrative analysis of RNA and mC features cross-validates neuronal cell clusters. (**A**) Schematic diagram of the cluster cross-validation strategy using matched single-cell methylome and transcriptome profiles. (**B-C**) Mean squared error (MSE) between RNA expression profile (B) or mCH (C) of individual cells and cluster centroids were plotted as a function of the number of clusters. The points of minimum MSE and minimum MSE + 1 standard error are indicated by arrows. Cross-validation analysis was performed in reciprocal directions by performing Leiden clustering using mC (B) or RNA (C) profiles followed by cross-validation using the matched RNA (B) and mC (C) data, respectively. (**D**) Schematic diagram of the over- and under-splitting analysis using matched single-cell methylome and transcriptome profiles. (**E**) Over-splitting of mC-defined clusters were quantified by the fraction of cross-modal *k*-partners found in the same cluster defined by RNA. Shades indicate confidence intervals of the mean. (**F**) Under-splitting of clusters was quantified as the cumulative distribution function of normalized self-radius. (**G**) Scatter plot of over-splitting (S_over_) and under-splitting (S_under_) scores for all neuronal clusters. Dot sizes represent cluster size. The actual data trend shows a linearly regressed line on both major clusters and sub clusters. (**H**) Joint UMAP visualization of snmC2T-seq transcriptome and methylome by computational integration using the SingleCellFusion method, assuming snmC2T-Seq transcriptomes and methylome were derived from independent datasets. (**I**) Accuracy of computational integration determined by the fraction of cells with matched transcriptome and epigenome profile grouped in the same cluster. (**J**) Confusion matrix normalized by each row. Each row shows the fraction of cells from each joint cluster that are from each cluster defined in Fig 2. Transcriptomes and DNA methylomes are quantified separately.

We further developed metrics to quantify over-splitting and under-splitting (Figure 4D, Figure S4B) by defining a graph connecting each cell to *k* cells with the greatest cross-modality similarity (called *k*-partners). An over-splitting score was calculated as the fraction of each cell’s *k*-partners which are in the same cluster (Methods; Figure 4D,E). This measure shows that major clusters resemble ideal, homogeneous clusters (simulated by shuffling gene features) and have little over-splitting (Figure 4E, Figure S4C). Most sub-clusters also had relatively little over-splitting, though some were more distinct than others (Figure S4C,E). To quantify under-splitting, we reasoned that all cells in a cluster should be statistically equivalent. Therefore, each cell’s mC profile should be no more correlated with its own RNA profile than with the RNA profile of any other cell of the same type. This could be violated if there is any discrete or continuous variation within the cluster that is correlated between modalities. Using a cross-modal score based on each cell’s cross-modal self-radius (see Methods), we found that major neuronal types had substantial within-cluster variation across cells indicating under-splitting (Figure 4F, Figure S4D,F). By contrast, subtypes resembled ideal (shuffled) clusters to a greater degree. Combining both scores, we could quantitatively map the lumper-splitter tradeoff in terms of the degree of over- and under-splitting for each major type or subtype (Figure 4G).

Integration of single-cell genomics data has been a focus of recent computational studies, yet existing methods lack validation on ground truth from experimental single-cell multi-omic dataset (Stuart and Satija, 2019). By treating snmC2T-seq transcriptome and mC profiles as if they were generated from different single cells, we could test the performance of computational integration using LIGER (Welch et al., 2019) Figure S4G (Welch et al., 2019) and Single Cell Fusion (Methods; Figure 4H). We quantified the accuracy of computational integration as the fraction of cells whose transcriptome and mC profiles were assigned to the same cluster (Figure 4I, Figure S4H-I). Both methods integrated the two data modalities well at the major neuronal type level, achieving an overall accuracy of 87.3% (Figure 4I-J). As expected, computational integration of fine-grain cell subtypes was less accurate (62.6%) and more variable across clusters (Figure 4I), potentially because of the greater degree of over-clustering (Figure 4E).

### Diverse correlation between gene body mCH and gene expression

Using the paired profiling of transcriptome and mC by snmC2T-Seq, we found diverse patterns of correlation between mCH and gene expression across thousands of single cells. Figure 5A shows examples for three distinct types of correlations between gene body mCH and gene expression. *KCNIP4* shows inverse correlation between mCH and RNA across a broad range of cell types. *ADARB2* is a marker gene for CGE-derived inhibitory cells and showed strong inter-cluster correlation, but no intra-cluster correlation between mCH and RNA. Finally, *GPC5* has a gradient of mCH across clusters (low in CGE VIP, high in L1-3 CUX2), but no corresponding pattern of differential gene expression across cell types. Applying this correlation analysis to all 13,637 sufficiently covered genes, we found that 38% (n=5,145) have significant negative correlation between mCH and RNA (mCH-RNA coupled, FDR < 5%). The majority of genes (62%) had no apparent correlation that could be distinguished from noise (mCH-RNA uncoupled, Figure 5B). The pattern of correlation was highly consistent between the specimens we profiled, and robust with respect to normalization (Figure S5A-F). We found that genes with significant correlation between mCH and gene expression are longer, more highly expressed, and are enriched in neuronal functions (Figure S5G-I).

**Figure 5.**
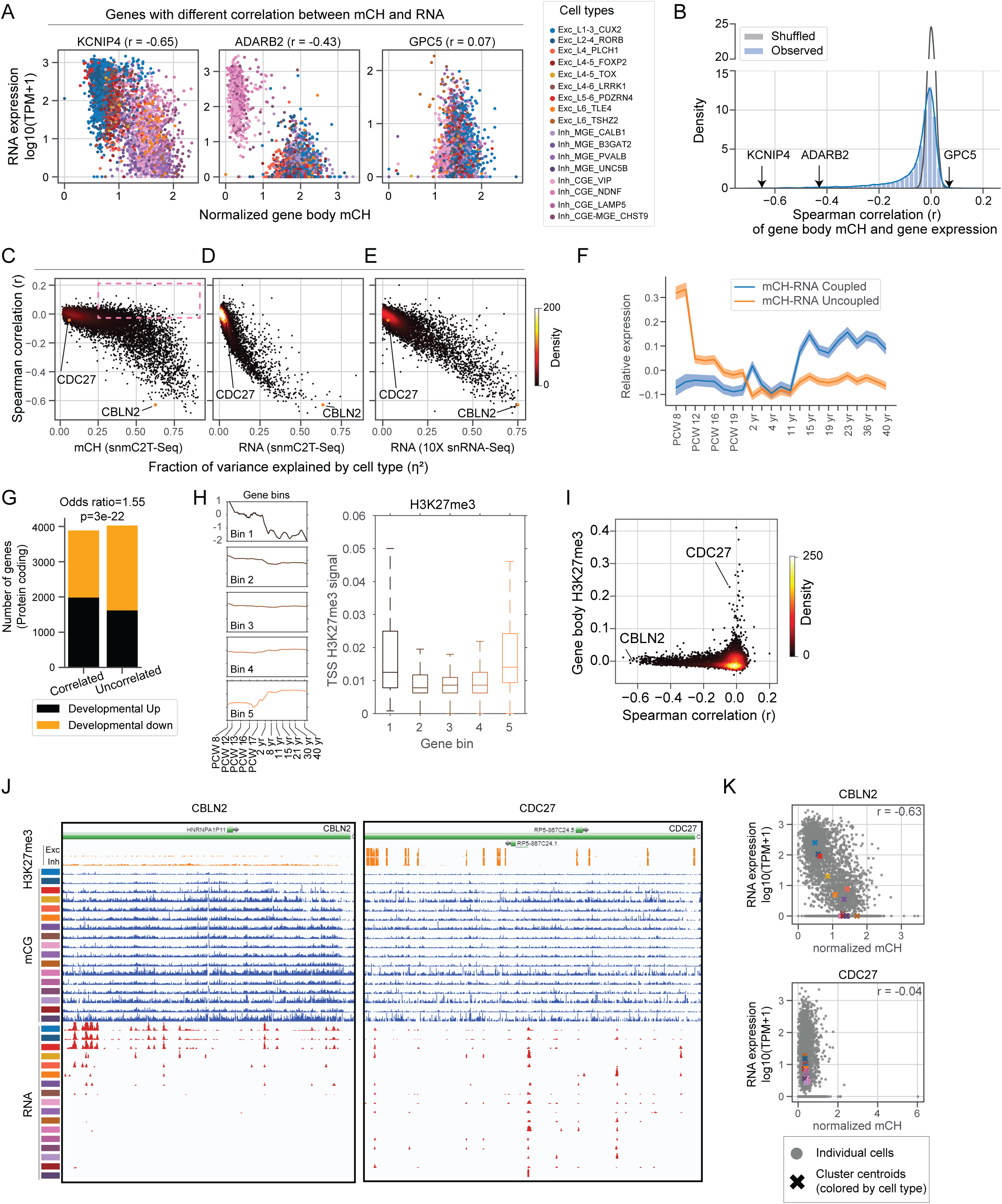
Single-cell correlation analysis of RNA expression and gene body non-CG methylation. (**A**) Scatter plots of gene body mCH (normalized by the global mean mCH of each cell) and gene expression (log_10_(TPM+1)) of example genes (KCNIP4, ADARB2, GPC5) across all neuronal cells. Cells are colored by major cell types defined in Figure 2. The spearman correlation coefficient (r) is shown for each example gene. (**B**) Distribution of Spearman correlation coefficient between gene expression and gene body mCH. Blue represents the actual distribution; Gray represents the distribution with randomly shuffled cell labels. (**C-E**) Scatter plot of correlation coefficient of gene body mCH and RNA versus the fraction of variance explained by cell type (η^2^) from 3 different datasets/features: snmC2T-Seq mCH, snmC2T-Seq RNA, and snRNA-Seq. (**F)** Line plot of mean relative expression over developmental time points with 2 different gene groups (mCH-RNA coupled in blue; mCH-RNA uncoupled in orange). Relative expression level is defined as the log2(RPKM) minus mean log2(RPKM) over all time points for each gene. (**G**) Barplot of the number of genes (protein-coding) in each of the 4 categories according to whether it’s developmentally up- or down-regulated, and whether its mCH-RNA is coupled or not. (**H**) Left: Line plots of mean relative expression level over developmental time points for 5 gene bins. Genes are binned by gene expression ratio between early fetal (PCW 8-9) and adult (>2 yrs). Right: boxplot of TSS H3K27me3 signals at each of the 5 gene bins. (**I**) Scatter plot of Spearman correlation of gene body mCH and gene expression versus the mean H3K27me3 signal in neurons at gene body level. The H3K27me3 ChIP-seq data is from purified Glutamatergic and GABAergic neurons from human frontal cortex (Kozlenkov et al, 2018). (**J**) Genome browser track visualization of CBLN2 and CDC27. (**K**) Gene level signal of CBLN2 and CDC27: scatter plot of normalized gene body mCH versus gene expression for all neuronal cells. Raw mCH level is normalized by the global mean mCH level of each cell.

We further investigated the factors that determine the degree of correlation between mCH and RNA for each gene. We reasoned that housekeeping genes with strong expression and little variation across cell types would show weak mCH-RNA correlation, whereas mCH-RNA coupling is enriched in genes with cell-type specific expression. We quantified the cell-type specificity of gene expression and DNA methylation by calculating the fraction of variance in gene expression explained by cell type (*RNA* η^2^ and *mCH* η^2^, Figure 5C-E and Figure S5J). Consistent with our hypothesis, genes with greater *RNA* η^2^ had a stronger inverse-correlation between mCH and RNA (Figure 5D-E). Notably, we found a large number of genes (n=1,243) with strong gene body mCH diversity across cell types (*mCH* η^2^ > 0.25) but no apparent correlation between mCH and RNA (r < −0.03) (box in Figure 5C). This suggests that lack of correlation between mCH and gene expression is driven by variability in gene expression within cell types despite conserved DNA methylation signatures.

The accumulation of mCH in the frontal cortex starts from the second trimester of embryonic development and continues into adolescence (Lister et al., 2013; Luo et al., 2016). The developmental dynamics of mCH motivated us to compare the developmental expression of mCH-RNA coupled and uncoupled genes. We found that mCH-RNA uncoupled genes, on average, are highly expressed during early fetal brain development (PCW 8-9) and are later repressed, whereas the expression of mCH-RNA coupled genes are moderately increased during development (Figure 5F). Consistently, developmentally down-regulated genes are significantly enriched in the mCH-RNA uncoupled group (Figure 5G). We speculated that the developmentally down-regulated genes may be repressive by alternative epigenomic marks such as histone H3K27 trimethylation (H3K27me3), which leads to the uncoupling of RNA and gene body mCH. By binning all the genes by their expression dynamics during brain development, we indeed found the promoter of both down- and up-regulated genes are enriched in H3K27me3 and depleted in active histone marks (Figure 5H and Figure S5K). We directly compared mCH-RNA correlation and H3K27me3 in purified human cortical glutamatergic and GABAergic neurons (Kozlenkov et al., 2018), and found genes with strong H3K27me3 signal clearly show weak correlations between gene body mCH and gene expression (e.g. CDC27 Figure 5I-K). In summary, although mCH and gene expression are clearly inversely correlated at a global scale, substantial variations can be observed from genes to genes at a single-cell level and can be partially explained by the presence of alternative epigenetic pathways such as polycomb repression.

### DNA methylation signatures of hierarchical transcription factor regulation in neural lineages

Timed expression of transcription factors (TF) during specific developmental stages is critical for neuronal differentiation (Deneris and Hobert, 2014; Kepecs and Fishell, 2014). We hypothesized that the cell-type hierarchy reconstructed from mC information reflects the developmental lineage of human cortical neurons. If so, then key transcription factors that specify neuronal lineage can be identified for each branch of the hierarchy. We separately constructed hierarchies for inhibitory and excitatory neurons based on the concatenated principal components of mCH and mCG (Figure 6A and S6A). The inhibitory neuron hierarchy comprises two major branches corresponding to medial ganglionic eminence (MGE) and caudal ganglionic eminence (CGE) derived cells. These major populations contain intermediate neuronal populations such as PVALB-expressing Basket Cell (BC) and Chandelier Cell (ChC), or the recently reported LAMP5-expressing Rosehip neurons (Boldog et al., 2018). At the finest level, the hierarchy contains 33 neuronal subtypes (Figure 6A). To identify TFs involved in the specification of neuronal lineages, we compared three levels of molecular information for each of 1639 human TFs (Lambert et al., 2018) between the daughter branches (Figure 6B). To assess the genome-wide DNA binding activity of the TF at regulatory elements, we used enrichment of DNA binding sequence binding motifs in differentially methylated regions (DMRs). To assess TF gene expression, we used both mRNA expression and TF gene body mCH level.

**Figure 6.**
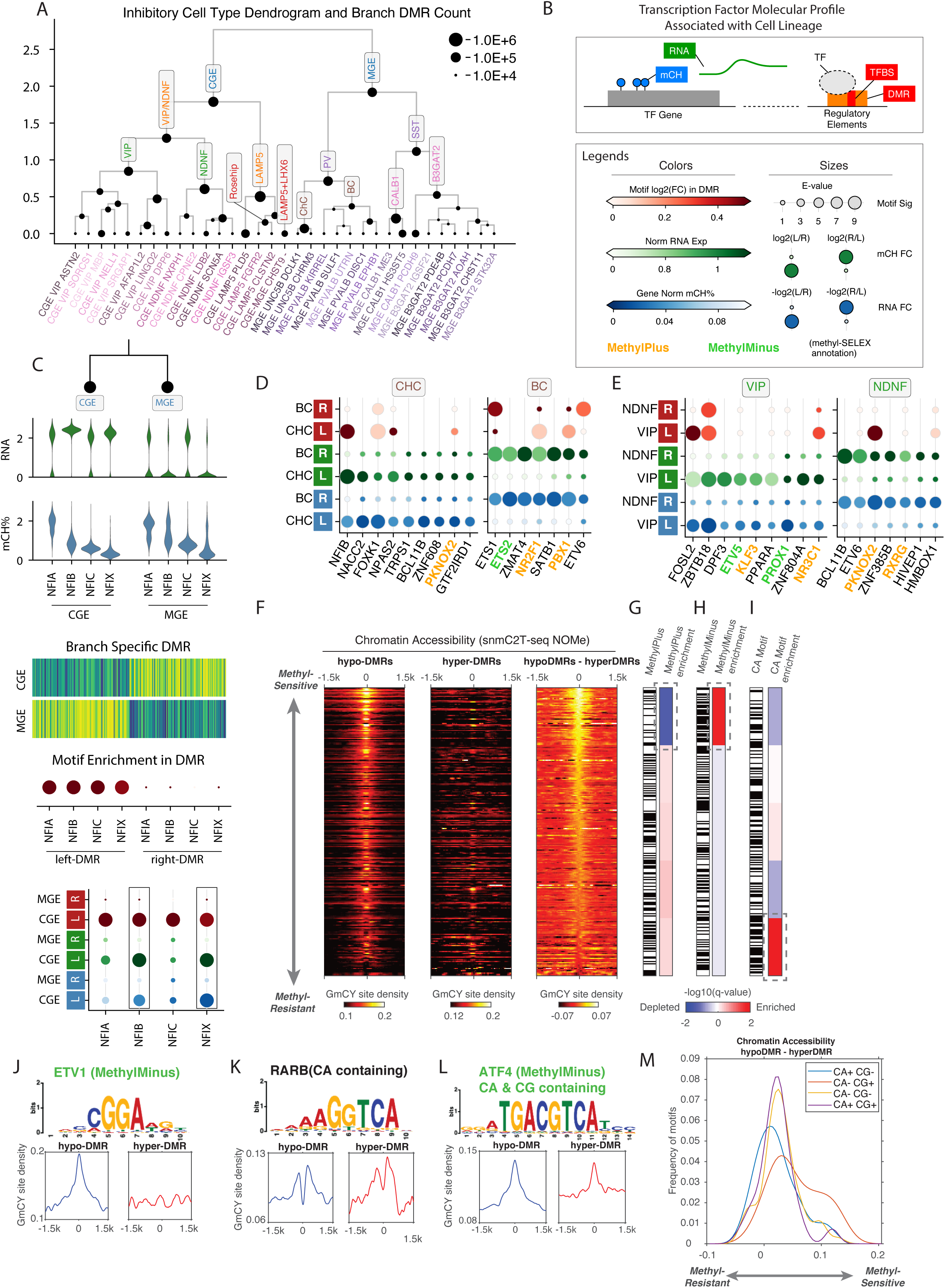
DMR phylogeny and transcription factor hierarchy in the human cortex. (**A**) Inhibitory neuron subtype dendrogram. The node size represents the number of DMRs detected between the left and right branches. Nodes corresponding to known inhibitory cell type groups are annotated in the dendrogram. (**B**) Schematics of the three levels of molecular information we use to identify candidate TF related to the specific lineage. (**C**) The workflow of TF analysis using the NFI family as an example. Three kinds of information are gathered for each of the TF genes: 1. RNA expression; 2. Gene body mCH level; 3. TF motif enrichment in the branch-specific DMR. We create a combined dot plot view for all three kinds of information, the genes show lineage specificity in both 1 and 2 are circled by black boxes. (**D-E**) The combined dot plot view for TFs showing ChC vs. BC (D) or VIP vs. NDNF (E) specificity in motif enrichment, RNA or mCH levels. Colors for every two rows from bottom to top: TF motif enrichment log2(fold change), branch mean expression log(1 + CPM), lineage mean gene body mCH level. Sizes for every two rows from bottom to top: E-value of the motif enrichment test, relative fold change of expression level, relative fold change of mCH level between the two branches. Colors for the motif names: TF motif methylation preference annotated by methyl-SELEX experiment (Yin et al., 2017), orange indicate MethylPlus, green indicate MethylMinus. (**F**) The binding of TFs to hypermethylated regions validated by chromatin accessibility measurement using the snmC2T-seq NOMe-seq profile. (**G-I**) Enrichment or depletion of MethylPlus TFs (G), MethyMinus TFs (H) and TFs whose binding motif containing CA dinucleotides (I). (**J-L**) Examples of chromatin accessibility profiles at the binding motifs of ETV1 (MethylMinus) (J), RARB (motif contains CA) (K) and ATF4 (motif contains CA and CG) (L). (**M**) Comparison of the chromatin accessibility at the binding motifs containing CA or CG dinucleotides.

Our integrated strategy taking advantage of matched information for TF motif enrichment, transcript abundance and TF gene body mCH level allowed us to distinguish the relative importance of closely related TFs sharing a common binding motif based on their cell type-specific expression (Castro-Mondragon et al., 2017) (Figure 6C). For example, we predicted that NFIB and NFIX contribute to CGE lineage specification since they show greater RNA abundance and stronger gene body mCH depletion than closely related TFs NFIA and NFIC. We systematically applied this approach across the inhibitory neuron hierarchy, using 579 curated motifs from the JASPAR 2018 CORE vertebrates database (Figure S6B-I) (Fornes et al., 2019). Many predicted lineage regulators were homologous to cell-type lineage regulators in mouse cortical development, such as NFIX, NFIB for CGE-derived neurons (Figure S6F), or LHX6, SOX6 and SATB1 for MGE-derived neurons (Figure S6F) (Kepecs and Fishell, 2014; Paul et al., 2017). The motifs of some TFs were also recurrently enriched in multiple lineages. For example, the NFIB gene (Piper et al., 2014) is not only specific to CGE neurons but also highly expressed and hypo-methylated in PV-expressing chandelier cell (ChC) but not basket cells (BC) (Figure 6D). The same expression pattern of NFIB was found in a comparison of mouse ChC - BC (Paul et al., 2017). These findings provide cogent evidence that the conserved major cell types of human and mouse (Hodge et al., 2019) also have shared basic rules of TF regulation. The same TF gene may perform multiple roles in different cell type lineages.

Previous studies including ours have found that discrete genomic regions with reduced mCG (hypomethylated DMRs) mark active regulatory elements (Mo et al., 2015; Schultz et al., 2015; Stadler et al., 2011; Ziller et al., 2013). We expected that TF binding motifs would be enriched in hypomethylated DMRs for cell types where the TF gene is active expressed and has low gene-body mCH. However, we identified several TFs with an opposite pattern: their binding motif was enriched in the hypomethylated DMRs of the alternative lineage showing low TF expression and high gene body mCH. For example, the motifs of NR2F1 and PBX1 were enriched in the hypomethylated DMRs of ChC, but both TFs were actively expressed in BC and not ChC (Figure 6D). Similarly, the PKNOX2 motif was enriched in hypomethylated DMRs of VIP cells, yet *PKNOX2* is preferentially expressed in NDNF neurons (Figure 6E). These data suggest that certain TFs can preferentially bind to hypermethylated regions (i.e. hypomethylated regions in the alternative lineage). This non-classical preference for methylated binding sites has been demonstrated in previous *in vitro* studies (Hu et al., 2013; Yin et al., 2017). In particular, Yin et al. used an *in vitro* assay to bind each recombinant TF protein to a pool of synthetic DNA (methyl-SELEX). They identified hundreds of TFs whose binding is inhibited (MethylMinus) or promoted (MethylPlus) by the presence of methylated CpG sites in their binding motifs. We analyzed the *in vivo* binding of MethylPlus TFs to hypermethylated DNA by analyzing chromatin accessibility measured by the snmC2T-seq NOMe-seq profile (Figure 6F). We quantified the average chromatin accessibility at TF binding motifs that are lowly methylated (overlapping with hypomethylated DMRs) or highly methylated (overlapping with hypermethylated DMRs) (Figure 6F), and used the difference in chromatin accessibility to determine the *in vivo* sensitivity of each TF to cytosine methylation. We found a general agreement between our *in vivo* approach and the *in vitro* methyl-SELEX results with MethylMinus TFs showing enrichment in the upper part of Figure 6F (e.g. ETV1 in Figure 6J), which showed greater chromatin accessibility between lowly and highly methylated TF motifs (Figure 6H). Consistently, MethylPlus TFs are strongly depleted in the upper part of Figure 6F (Figure 6G). Therefore, our joint analysis of mC and chromatin accessibility using snmC2T-seq provided *in vivo* evidence for the modulation of TF binding by cytosine methylation.

Lastly, we examined the correlation between chromatin accessibility and the presence of CA dinucleotide in the TF binding motifs, since CA is the predominant sequence context of mCH in the human brain (Lister et al., 2013). Intriguingly, we found a significant enrichment of TF binding motifs containing CA dinucleotides in the lowest part of Figure 6F (Figure 6I), suggesting the accessibility of TF binding motifs containing CA is less affected by mC. Across all TF binding motifs examined, the accessibility of motifs containing both CA and CG dinucleotides (CA+ CG+, p-value = 1 x 10^-4^, e.g. ATF4, Figure 6L) or only CA (CA+ CG-, p-value = 5.7 x 10^-6^, e.g. RARB, Figure 6K) show significantly less sensitivity to mC than motifs containing CG dinucleotides only (CA-CG+) (Figure 6M). The results suggest certain TFs may be able to bind hypermethylated regions through the interaction with mCA sites. The modulation of TF binding by mCA has not been systematically explored since previous studies have focused on the effect of mCG sites (Hu et al., 2013; Yin et al., 2017).

### Cortical cell regulatory genomes predict developmental and adult cell types involved in neuropsychiatric diseases

The strong enrichment of disease heritability in gene regulatory elements has allowed the prediction of disease-relevant cell types using epigenomic signatures (Finucane et al., 2015), including neuropsychiatric disorders (Kozlenkov et al., 2018). By reconstructing mC and open chromatin maps from single-cell profiles, we used LD score regression partitioned heritability to infer the relevant cell types for a set of neuropsychiatric traits using DMRs and ATAC-seq peaks (Table S9-10) (Finucane et al., 2015). To capture regulatory elements active during early development which may implicated in psychiatric disease, we further included DMRs identified from bulk fetal (PCW 19) human cortex methylome (Luo et al., 2016) and DNase-seq peaks identified from fetal brain samples (Roadmap Epigenomics Consortium et al., 2015). Using a statistical threshold of FDR < 0.1, we identified 27 disease-cell type associations across 23 cortical cell types or bulk samples for 16 neuropsychiatric traits (Figure S7A). Each association corresponds to a significant enrichment of disease heritability within the corresponding cell type’s active regulatory regions. By contrast, only 3 associations were found in DMRs identified from 18 bulk non-brain tissues (Figure S7A) (Schultz et al., 2015). This result strongly suggests our partitioned heritability analysis has correctly identified the brain as the relevant tissue types for neuropsychiatric traits.

Using our single-cell epigenomic dataset, we identified enrichment of heritability in distinct cortical cell types for a number of psychiatric disorders. In most cases our analysis enhanced the cell-type resolution of partitioned heritability analysis compared with previous efforts. For example, using single-cell RNA-seq dataset, the genetic risk of schizophrenia (SCZ) was previously mapped to broad cortical neuronal populations including neocortical somatosensory pyramidal cells, and cortical interneurons (Kozlenkov et al., 2018; Skene et al., 2018). Our analysis further identified the enrichment of SCZ heritability in multiple types of intratelencephalic (IT) neuron types (L1-3 CUX2, L4-5 FOXP2 and L5-6 PDZEN4), in addition to a medial ganglionic eminence (MGE) derived inhibitory cell type (MGE CALB1) (Figure 7A). Intriguingly, the heritability of bipolar disorder (BP) was specifically enriched in a deep layer IT neuron type L5-6 PDZEN4 (Figure 7B). We also found a specific enrichment of autism spectrum disorder (ASD) risk in a deep-layer thalamic-projecting neuronal population L6 TLE4 (Figure 7C). By contrast, the heritability of educational attainment (EA) was broadly distributed across multiple types of neurons including excitatory cells (L1-3 CUX2 and L4 PLCH1) and inhibitory neurons derived from both CGE (CGE LAMP5) and medial ganglionic eminence (MGE, MGE CALB1) (Figure 7D). Consistent with the neurodevelopmental hypothesis that gene mis-regulation during brain development underlies many psychiatric disorders (Birnbaum and Weinberger, 2017), fetal cortex DMRs are enriched in the heritability for 9 neuropsychiatric traits including SCZ and educational attainment (Figure S7A). However, the partitioned heritability analysis using the fetal cortex sample is likely underpowered due to the cell-type heterogeneity.

**Figure 7.**
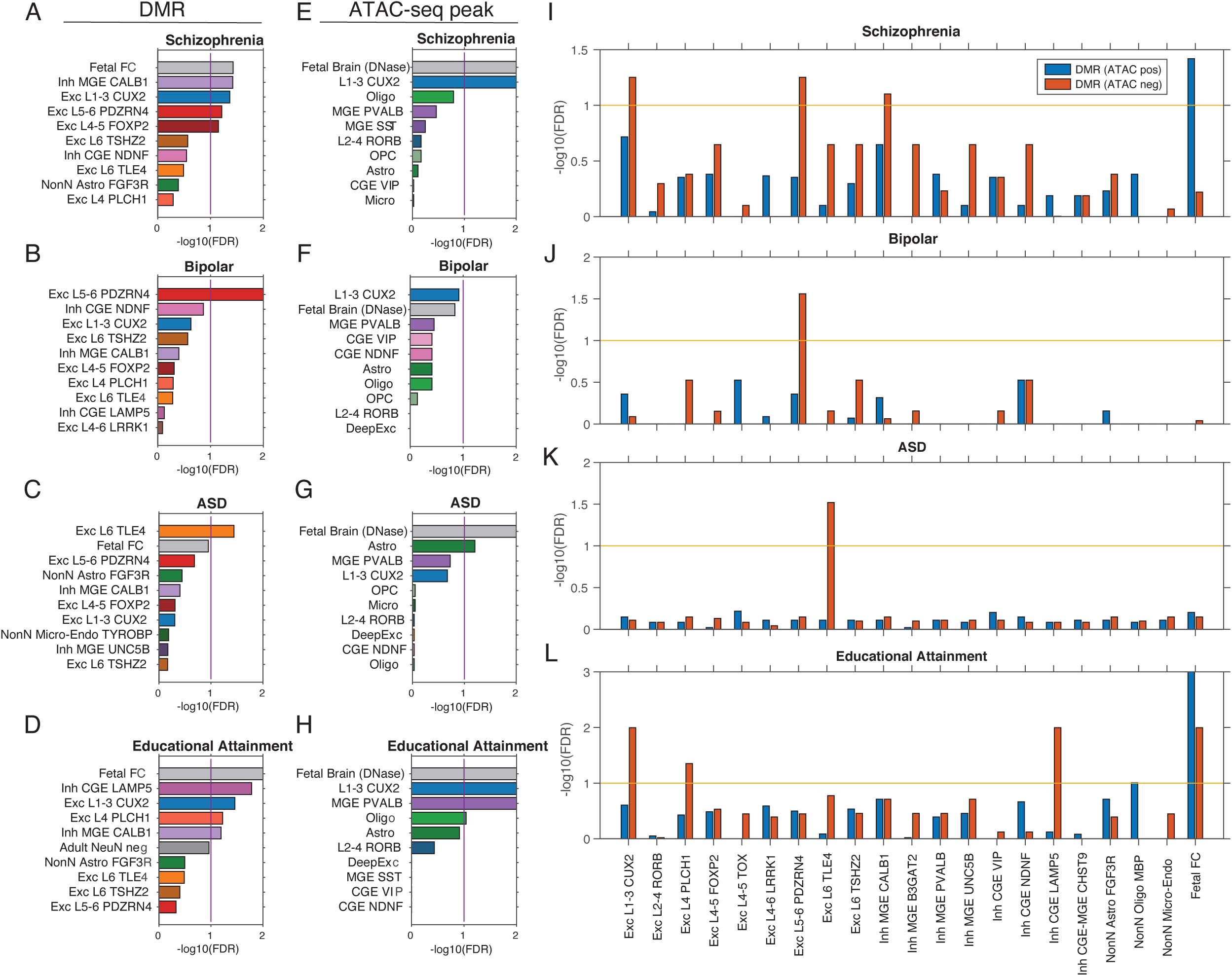
Identification of brain cell types involved in neuropsychiatric traits. (**A-H**) Partitioned heritability analysis using cell-type specific DMRs (A-D) or ATAC-seq peaks (E-H). (**I-L**) Partitioned heritability analysis using DMRs stratified for the overlap with open chromatin regions.

We performed the partitioned heritability analysis using the two complementary types of epigenomic signatures: DMRs and open chromatin regions (ATAC-seq or DNase-seq peaks). To our surprise, the results obtained using DMRs and ATAC-seq peaks were substantially different. For example, the partition of SCZ heritability across DMRs identified enrichment in four adult cell types in addition to fetal cortex (Figure 7A), whereas the analysis using open chromatin regions only found enrichment in L1-3 CUX2 cells and the fetal brain (Figure 7E). To understand this discrepancy, we stratified DMR regions into two groups [DMR (ATAC-pos) and DMR (ATAC-neg)] by their overlap with open chromatin regions. Partitioned heritability across the stratified DMR regions revealed that in adult cells, DMR regions without open chromatin signature are more strongly enriched in heritability for SCZ, BP, ASD and EA. In fetal cortex, however, a stronger enrichment of SCZ and EA heritability was found in DMRs associated with open chromatin.

We speculate that DMRs without open chromatin contain vestigial enhancers (Hon et al., 2013), which contribute to the enrichment of disease heritability. Vestigial enhancers are active regulatory elements during embryonic development but become dormant in adult tissues (Hon et al., 2013). However, vestigial enhancers remain lowly methylated in adult tissues and can be identified as DMRs. Thus, vestigial enhancers can be strongly enriched in the genetic risk of neuropsychiatric traits since these regions are active regulatory elements during brain development. We identified the fraction of adult brain DMRs that correspond to vestigial enhancers, i.e. overlapping with open chromatin regions in the embryonic, but not the adult brain. Consistent with our speculation, in many cases vestigial enhancers show stronger enrichment of disease heritability (Figure S7I-L). In particular, the enrichment of ASD genetic risk in L4 PLCH1 and L6 TLE4 cells can only be identified in vestigial enhancers (Figure S7K). In summary, we found that single cell type DMRs integrates regulatory information in both developmental and adult brain and can be effectively used to predict cell types involved in neuropsychiatric disorders.

## DISCUSSION

Epigenomic studies often incorporate multiple molecular profiles from the same sample to explore possible correlations between gene regulatory elements and expression. The need for multi-omic comparison poses a challenge for single-cell analysis, since most existing single-cell techniques terminally consume the cell, precluding multi-dimensional analysis. To address this challenge, we have developed two single-nucleus multi-omic assays snmCT-seq and snmC2T-seq to jointly profile the transcriptome, DNA methylome and chromatin accessibility and can be applied to either single cells or nuclei from frozen human tissues. snmC2T-seq requires no physical separation of DNA and RNA and is designed to be a “single-tube” reaction for steps before bisulfite conversion to minimize material loss. snmC2T-seq is fully compatible with high-throughput single-cell methylome techniques such as snmC-seq2 (Luo et al., 2018) and can be readily scaled to analyze thousands of cells/nuclei.

The continuous development of multi-omic profiling techniques such as scNMT-seq (Clark et al., 2018) and snmC2T-seq, and several methods for joint RNA and chromatin accessibility profiling sci-CAR (Cao et al., 2018), SNARE-seq (Chen et al., 2019) and Paired-seq (Zhu et al., 2019) provide the opportunity to classify cell types with multiple molecular signatures. Our study provided a computational framework to cross-validate clustering-based cell-type classifications using multi-modal data. Through cross-validation between matched single-cell mC and RNA profiles, we found that between 20-50 human cortical cell types can be identified from our moderate size snmC2T-seq dataset (4,358 cells) with sound cluster robustness. This is consistent with the number of human frontal cortex cell types we reported in our previous (21 major types, (Luo et al., 2017)) and current (20 major types and 63 subtypes) studies. Using snmC2T-seq as a “ground-truth”, we determined that computational multi-modal integration tools perform well at the major cell-type level but show variable accuracy for the integration of fine grain subtypes. The computational strategies developed in this study can be applied to other types of multi-omic profiling including methods involving physiological measurement such as Patch-seq (Cadwell et al., 2016; Fuzik et al., 2016).

Epigenomic studies at both bulk and single-cell levels have established both mC and open chromatin as reliable markers for regulatory elements (Kelsey et al., 2017). However, the difference between the information provided by the two epigenomic marks has been less clear in the context of normal development and diseases. Our study found that DMRs contain disease-related regulatory information of both adult and embryonic tissues, with vestigial enhancers (Hon et al., 2013) as a possible mechanism that informs developmental gene regulation. The strong enrichment of genetic risks for neuropsychiatric disorders in vestigial enhancers enabled the prediction of causal cellular lineage for diseases using DMRs for partitioned heritability analyses and identified more diverse disease-relevant brain cell populations than similar analyses using open chromatin regions. The abundance of developmental information in DNA methylome suggests the possibility to study developmental processes and gene regulation in cell lineages using methylome profiling of adult tissues, especially given the practical and ethical challenges for obtaining primary human tissues from developmental stages.

## ACKNOWLEDGEMENTS

This work was supported by NIH grants: 5R21HG009274, 5R21MH112161 and 5U19MH114831 to J.R.E; U01MH114812 to E.L. J.R.E. is an Investigator of the Howard Hughes Medical Institute. W.D. is supported by a NIH training award 5T32MH020002. Postmortem human brain tissues were obtained from the NIH NeuroBioBank at the University of Maryland Brain and Tissue Bank and the University of Miami Brain Endowment Bank. We thank the donors and their families for their invaluable donations for the advancement of science. We would like to thank the QB3 Macrolab at UC Berkeley for purification of Tn5 transposase. Work at the Center for Epigenomics was supported in part by the UC San Diego School of Medicine.

## AUTHOR CONTRIBUTIONS

J.R.E, and C.L. conceived the study. J.R.E, E.A.M, M.M.B, B.R., E.L. S.L. and J.R.D supervised the study. C.L., B-A.W. and Z.Z. developed the scmCT-seq, snmCT-seq and snmC2T-seq methods. C.L., B-A.W., R.C., A.B., A.R., and J.R.N. generated the scmCT-seq, snmCT-seq and snmC2T-seq data. C.L., R.C. and J.R.N. generated the snmC-seq data. K.S., T.E.B., R.D.H, L.H., S.L. and E.L. generated and analyzed the snRNA-seq data. R.F., S.P., X.W. and B.R. generated and analyzed the snATAC-seq data. D.A.D. and D.C.M. acquired human brain specimen. D-S.L. and J.R.D reanalyzed the sn-m3C-seq data. H.L., F.X., C.L., W.D., E.J.A., D-S.L., J.Z., S-Y.N. analyzed the data. C.L., H.L. and F.X. drafted the manuscript. J.R.E, E.A.M, T.E.B., R.D.H, D.A.D and D.C.M edited the manuscript.

## Data Availability

Raw and processed data included in this study were deposited to NCBI GEO/SRA with accession number GSE140493. Methylome and transcriptomic profiles generated by scmCT-seq and snmCT-seq from H1 and HEK293T cells can be visualized at [http://neomorph.salk.edu/Human_cells_snmCT-seq.php]. snmC2T-seq generated from brain tissues can be visualized at [http://neomorph.salk.edu/human_frontal_cortex_ensemble.php].

## METHODS

### Cell cultures

HEK293T cells were cultured in DMEM with 15% FBS and 1% Penicillin-Streptomycin and dissociated with 1X TrypLE. H1 human ESCs (WA01, WiCell Research Institute) were maintained in feeder-free mTesR1 medium (Stemcell Technologies). hESCs (passage 26) were dispersed with 1U/ml Dispase and collected for single-cell sorting or nuclei isolation. For the sorting of single H1 and HEK293T cells, equal amounts of H1 and HEK293T cells were mixed and stained with anti-TRA-1-60 (Biolegend, Cat#330610) antibody.

### Human brain tissues

Postmortem human brain biospecimens GUID: NDARKD326LNK and NDARKJ183CYT were obtained from NIH NeuroBioBank at University of Miami Brain Endowment Bank. Postmortem human brain biospecimens UMB4540, UMB5577 and UMB5580 were obtained from NIH NeuroBioBank at University of Maryland Brain and Tissue Bank. Published snmC-seq was generated from frontal cortex (medial frontal gyrus) tissue obtained from a 25-year-old Caucasian male (UMB4540, labeled as M_25yr_1 in this study) with a postmortem interval (PMI) = 23 h. The snATAC-seq dataset was generated from specimen UMB4540. Additional snmC-seq data was generated in frontal cortex (superior frontal gyrus, Brodmann area 10) tissues obtained from a 58-year-old Caucasian male (GUID: NDARKD326LNK, labeled as M_58yr in this study) with a postmortem interval (PMI) = 23.4 h. snmC-seq2 data was generated from frontal cortex (Brodmann area 10) tissue from a 25-year-old Caucasian male (GUID: NDARKJ183CYT, labeled as M_25yr_2 in this study) with a PMI = 20.8 h. snmCT-seq and sn-m3C-seq data were generated from a 21-year-old Caucasian male (UMB5577, labeled as M_21yr in this study) with a PMI = 19h, and a 29-year-old Caucasian male (UMB5580, labeled as M_29yr in this study) with a PMI = 8h. The samples were taken from unaffected control subjects who died from accidental causes. The snRNA-seq dataset was generated from postmortem brain specimen H18.30.002 from the Allen Institute for Brain Science. Frontal cortex (BA44-45, 46) from this donor was used for the generation of single nucleus RNA-seq data. The donor was a 50 year old male with a PMI = 12 h.

### Nuclei isolation from cultured cells for scmCT-seq and snmCT-seq

Cell pellets containing 1 million cells were resuspended in 600 µl NIBT [250 mM Sucrose, 10 mM Tris-Cl pH=8, 25 mM KCl, 5mM MgCl_2_, 0.1% Triton X-100, 1mM DTT, 1:100 Proteinase inhibitor (Sigma-Aldrich P8340), 1:1000 SUPERaseIn RNase Inhibitor (ThermoFisher Scientific AM2694), 1:1000 RNaseOUT RNase Inhibitor (ThermoFisher Scientific 10777019)]. The lysate was transferred to a pre-chilled 2 ml dounce homogenizer (Sigma-Aldrich D8938) and dounced using loose and tight pestles for 20 times each. The lysate was then mixed with 400 µl of 50% Iodixanol (Sigma-Aldrich D1556) and gently pipetted on top of 500 µl 25% Iodixanol cushion. Nuclei were pelleted by centrifugation at 10,000 x g at 4°C for 20 min using a swing rotor. The pellet was resuspended in 2 ml of DPBS supplemented with 1:1000 SUPERaseIn RNase Inhibitor and 1:1000 RNaseOUT RNase Inhibitor. Hoechst 33342 was added to the sample to a final concentration of 1.25 nM and incubated on ice for 5 min for nuclei staining. Nuclei were pelleted by 1,000 x g at 4°C for 10 min and resuspended in 1 ml of DPBS supplemented with RNase inhibitors.

### Nuclei isolation from human brain tissues and GpC methyltransferase treatment for snmC2T-seq

Brain tissue samples were ground in liquid nitrogen with cold mortar and pestle, and then aliquoted and store at −80°C. Approximately 100mg of ground tissue was resuspended in 3 ml NIBT (250 mM Sucrose, 10 mM Tris-Cl pH=8, 25 mM KCl, 5mM MgCl_2_, 0.2% IGEPAL CA-630, 1mM DTT, 1:100 Proteinase inhibitor (Sigma-Aldrich P8340), 1:1000 SUPERaseIn RNase Inhibitor (ThermoFisher Scientific AM2694), 1:1000 RNaseOUT RNase Inhibitor (ThermoFisher Scientific 10777019)). The lysate was transferred to a pre-chilled 7 ml dounce homogenizer (Sigma-Aldrich D9063) and dounced using loose and tight pestles for 40 times each. The lysate was then mixed with 2 ml of 50% Iodixanol (Sigma-Aldrich D1556) to generate a nuclei suspension with 20% Iodixanol. Gently pipet 1 ml of the nuclei suspension on top of 500 µl 25% Iodixanol cushion in each of the 5 freshly prepared 2ml microcentrifuge tubes. Nuclei were pelleted by centrifugation at 10,000 x g at 4°C for 20 min using a swing rotor. The pellet was resuspended in 1ml of DPBS supplemented with 1:1000 SUPERaseIn RNase Inhibitor and 1:1000 RNaseOUT RNase Inhibitor. A 10 µl aliquot of the suspension was taken for nuclei counting using a Biorad TC20 Automated Cell Counter. One million nuclei aliquots were pelleted by 1,000 x g at 4°C for 10 min and resuspended in 200 µl of GpC methyltransferase M.CviPI (NEB M0227L) reaction containing 1X GC Reaction Buffer, 0.32 nM S-Adenoslylmethionime, 80U 4U/µl M.CviPI, 1:100 SUPERaseIn RNase Inhibitor and 1:100 RNaseOUT RNase Inhibitor and incubated at 37°C for 8 min. The reaction was stopped by adding 800 µl of ice-cold DPBS with 1:1000 RNase inhibitors and mixing. Hoechst 33342 was added to the sample to a final concentration of 1.25 nM and incubated on ice for 5 min for nuclei staining. Nuclei were pelleted by 1,000 x g at 4°C for 10 min, resuspended in 900 µl of DPBS supplemented with 1:1000 RNase inhibitors and 100 µl of 50mg/ml Ultrapure^TM^ BSA (Ambion AM2618) and incubated on ice for 5 min for blocking. Neuronal nuclei were labeled by adding 1 µl of AlexaFluor488-conjugated anti-NeuN antibody (clone A60, MilliporeSigma MAB377XMI) for 20 min.

### Reverse transcription for snmC2T-seq

Single cells or single nuclei were sorted into 384-well PCR plates (ThermoFisher 4483285) containing 1 µl mCT-seq reverse transcription reaction per well. The mCT-seq reverse transcription reaction contained 1X Superscript II First-Strand Buffer, 5mM DTT, 0.1% Triton X-100, 2.5 mM MgCl_2_, 500 µM each of 5’-methyl-dCTP (NEB N0356S), dATP, dTTP and dGTP, 1.2 µM dT30VN_4 oligo-dT primer (5’-AAGCAGUGGUAUCAACGCAGAGUACUTTTTTUTTTTTUTTTTTUTTTTTUTTTTTV N-3’ was used the cultured cell scmCT-seq and snmCT-seq experiments; 5’-/5SpC3/AAGCAGUGGUAUCAACGCAGAGUACUTTTTTUTTTTTUTTTTTUTTTTTU TTTTTVN-3’ was used for human brain snmC2T-seq experiments), 2.4 µM TSO_3 template switching oligo (/5SpC3/AAGCAGUGGUAUCAACGCAGAGUGAAUrGrG+G), 1U RNaseOUT RNase inhibitor, 0.5 U SUPERaseIn RNase inhibitor, 10U Superscript II Reverse Transcriptase (ThermoFisher 18064-071). For snmCT-seq and snmC2T-seq, the reaction further included 2 µM N6_2 random primer (/5SpC3/AAGCAGUGGUAUCAACGCAGAGUACNNNNNN). After sorting, the PCR plates were vortexed and centrifuged at 2000 x g. The plates were placed in a thermocycler and incubated using the following program: 25°C for 5 min, 42°C for 90min, 70°C 15min followed by 4°C.

### cDNA amplification for scmCT-seq, snmCT-seq and snmC2T-seq

3 µl of mCT-seq cDNA amplification mix was added into each mCT-seq reverse transcription reaction. mCT-seq cDNA amplification reaction containing 1X KAPA 2G Buffer A, 600 nM ISPCR23_2 PCR primer (/5SpC3/AAGCAGUGGUAUCAACGCAGAGU), 0.08U KAPA2G Robust HotStart DNA Polymerase (5 U/μL, Roche KK5517). PCR reactions were performed using a thermocycler with the following conditions: 95°C 3min -> [95°C 15 sec -> 60°C 30 sec -> 72°C 2min] -> 72°C 5min -> 4°C. The cycling steps were repeated for 12 cycles for scmCT-seq using H1 or HEK293 cells, 15 cycles for snmCT-seq using H1 or HEK293 cells and 14 cycles for snmC2T-seq using human brain tissues.

### Digestion of unincorporated DNA oligos for scmCT-seq, snmCT-seq and snmC2T-seq

For scmCT-seq and snmCT-seq using H1 and HEK293 cells, 1 µl uracil cleavage mix was added to into cDNA amplification reaction. Each 1 µl uracil cleavage mix contains 0.25 µl Uracil DNA Glycosylase (Enzymatics G5010) and 0.25 µl Endonuclease VIII (Enzymatics Y9080) and 0.5 µl Elution Buffer (Qiagen 19086). Unincorporated DNA oligos were digested at 37°C for 30 min using a thermocycler. We have found that Endonuclease VIII is dispensable for the digestion of unincorporated DNA oligos since the alkaline condition during the desulfonation step of bisulfite conversion can effective cleave abasic sites created by Uracil DNA Glycosylase (Greenberg, 2014). Therefore for snmC2T-seq using human brain tissues, each cDNA amplification reaction was treated with 1µl uracil cleavage mix containing 0.5 µl Uracil DNA Glycosylase (Enzymatics G5010-1140) and 0.5 µl Elution Buffer (Qiagen 19086).

### Bisulfite conversion and library preparation

Detailed methods for bisulfite conversion and library preparation are previously described for snmC-seq2 (Luo et al., 2017, 2018). The following modifications were made to accommodate the increased reaction volume of scmCT-seq, snmCT-seq or snmC2T-seq: Following the digestion of unused DNA oligos, 25 µl instead of 15 µl of CT conversion reagent was added to each well of a 384-well plate. 90 µl instead of 80 µl M-binding buffer was added to each well of 384-well DNA binding plate. scmCT-seq libraries were generated using the snmC-seq method as described in Luo et al., 2017 (Luo et al., 2017). snmCT-seq and snmC2T-seq libraries were generated using the snmC-seq2 method as described in Luo et al., 2018 (Luo et al., 2018). The scmCT-seq and snmCT-seq libraries were sequenced using an Illumina HiSeq 4000 instrument with 150 bp paired-end reads. The snmC2T-seq libraries generated from human brain specimens were sequenced using an Illumina Novaseq 6000 instrument with S4 flowcells and 150 bp paired-end mode.

### The mapping pipeline for snmC-seq, snmC-seq2, snmCT-seq, and snmC2T-seq

We implemented a versatile mapping pipeline (<u>cemba-data.rtfd.io</u>) for all the methylome based technologies developed by our group (Luo et al., 2017, 2018). The main steps of this pipeline include: 1) Demultiplexing FASTQ files into single-cell; 2) Reads level QC; 3) Mapping; 4) BAM file processing and QC; 5) final molecular profile generation. For snmC-seq and snmC-seq2, the details of the five steps are described previously (Luo et al., 2017, 2018). For scmCT-seq, snmCT-seq and snmC2T-seq, steps 1 and 2 are identical as snmC-seq2, steps 3 to 5 are split into “a” for methylome and “b” for transcriptome as following:

#### Step 3a (methylome)

To map methylome reads, reads from step 2 were mapped onto the human hg19 genome using bismark (Krueger and Andrews, 2011) with the same setting as snmC-seq2.

#### Step 3b (transcriptome)

To map transcriptome reads, reads from step 2 were mapped to GENCODE human v28 indexed hg19 genome using STAR 2.7.2b (Dobin et al., 2013) with the following parameters: *--alignEndsType EndToEnd --outSAMstrandField intronMotif --outSAMtype BAM Unsorted --outSAMunmapped Within --outSAMattributes NH HI AS NM MD --sjdbOverhang 100 --outFilterType BySJout --outFilterMultimapNmax 20 --alignSJoverhangMin 8 --alignSJDBoverhangMin 1 --outFilterMismatchNmax 999 --outFilterMismatchNoverLmax 0.04 --alignIntronMin 20 --alignIntronMax 1000000 --alignMatesGapMax 1000000 --outSAMattrRGline ID:4 PL:Illumina*.

#### Step 4a (methylome)

PCR duplicates were removed from mapped reads using Picard MarkDuplicates. The non-redundant reads were then filtered by MAPQ > 10. To select genomic reads from the filtered BAM, we used the “XM-tag” generated by bismark to calculate reads methylation level and keep reads with mCH ratio < 0.5 and the number of cytosines ≥ 3.

#### Step 4b (transcriptome)

the STAR mapped reads were first filtered by MAPQ > 10. To select RNA reads from the filtered BAM, we used the “MD” tag to calculate reads methylation level and keep reads with mCH ratio > 0.9 and the number of cytosines ≥ 3. The stringency of read partitioning was determined by applying the criteria for identifying snmCT-seq transcriptome reads to snmC-seq2 data (SRR6911760, SRR6911772, SRR6911776) (Luo et al., 2018), which contains no transcriptomic reads. Similarly, the criteria for identifying snmCT-seq methylome reads were applied to Smart-seq data (SRR944317, SRR944318, SRR944319, SRR944320) (Picelli et al., 2013), which contains no methylome reads.

#### Step 5a (methylome)

Tab-delimited (ALLC) files containing methylation level for every cytosine position was generated using methylpy *call_methylated_sites* function (Schultz et al., 2015) on the BAM file from the step 4a. For snmC2T-seq, an additional base was added before the cytosine in the context column of the ALLC file using the parameter “--num_upstr_bases 1”, to distinguish GpC sites from HpC sites for the NOMe-seq modality.

#### Step 5b (transcriptome)

BAM file from step 4b were counted across gene annotations using featureCount 1.6.4 (Liao et al., 2014) with the default parameters. Gene expression was quantified using either only exonic reads with “-t exon” or both exonic and intronic reads with “-t gene”.

### Methylome feature generation

After allc files were generated, the methylcytosine (*mc*) and total cytosine basecalls (*cov*) were summed up for each 100kb bin across the hg19 genome. For snmC-seq, snmC-seq2, and snmCT-seq, cytosine and methylcytosine basecalls in CH (H=A, T, C) and CG context were counted separately. For snmC2T-seq, the HCH context was counted for CH methylation and HCG is counted for CG methylation. The GCY (Y=T, C) context was counted as the chromatin accessibility signal (NOMe-seq in snmC2T-seq) and the HCY context was counted as the endogenous mCH background. In addition to the 100kb feature set, we also counted gene body methylation levels using gene annotation from GENCODE v28. The 100kb feature set was used in methylation-based clustering analysis and data integration; the gene body feature set was used in methyl-marker identification, cluster annotation and data integration between methylome and transcriptome.

#### Preprocessing of snmC-seq and snmC-seq2 data for clustering analyses Cell filtering

We filtered the cells based on these main mapping metrics: 1) mCCC rate < 0.03. mCCC rate reliably estimates the upper bound of bisulfite non-conversion rate (Luo et al., 2017), 2) overall mCG rate > 0.5, 3) overall mCH rate < 0.2, 4) total final reads > 500,000, 5) bismark mapping rate > 0.5. Other metrics such as genome coverage, PCR duplicates rate, index ratio were also generated and evaluated during filtering. However after removing outliers with the main metrics 1-5, few additional outliers can be found.

#### Feature filtering

100kb genomic bin features were filtered by removing bins with mean total cytosine base calls < 300 or > 3000. Regions overlap with the ENCODE blacklist (Amemiya et al., 2019) were also removed from further analysis.

#### Computation and normalization of the methylation rate

For CG and CH methylation, the computation of methylation rate from the methylcytosine and total cytosine matrices contains two steps: 1) prior estimation for the beta-binomial distribution and 2) posterior rate calculation and normalization per cell.

Step 1, for each cell we calculated the sample mean, *m*, and variance, *v*, of the raw mc rate *(mc / cov)* for each sequence context (CG, CH). The shape parameters (α, β) of the beta distribution were then estimated using the method of moments:

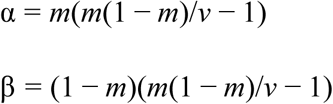

This approach used different priors for different methylation types for each cell, and used weaker prior on cells with more information (higher raw variance).

Step 2, We then calculated the posterior: 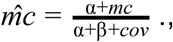 We normalized this rate by the cell’s global mean methylation, *m* = α/(α + β). Thus, all the posterior 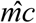 with 0 *cov* will be constant 1 after normalization. The resulting normalized *mc* rate matrix contains no NA (not available) value and features with less *cov* tend to have a mean value close to 1.

#### Selection of highly variable features

Highly variable methylation features were selected based on a modified approach using the scanpy package *scanpy.pp.highly_variable_genes* function (Wolf et al., 2018). In brief, the *scanpy.pp.highly_variable_genes* function normalized the dispersion of a gene by scaling with the mean and standard deviation of the dispersions for genes falling into a given bin for mean expression of genes. In our modified approach, we reasoned that both the mean methylation level and the mean *cov* of a feature (100kb bin or gene) can impact *mc* rate dispersion. We grouped features that falling into a combined bin of mean and *cov*, and then normalized the dispersion within each mean-*cov* group. After dispersion normalization, we selected the top 3000 features based on normalized dispersion for clustering analysis.

#### Dimension reduction and combination of different mC types

For each selected feature, *mc* rates were scaled to unit variance and zero mean. PCA was then performed on the scaled *mc* rate matrix. The number of significant PCs was selected by inspecting the variance ratio of each PC using the elbow method. The CH and CG PCs were then concatenated together for further analysis in clustering and manifold learning.

### Preprocessing of snmC2T-seq data for clustering analysis

#### Methylome preprocessing

The methylome modality preprocessing is similar to snmC-seq2 with one major modification: non-CG methylation is quantified using the HCH context; CG methylation is quantified using the HCG context. Chromosome 100kb bin features with mean total cytosine base calls between 250 and 2500 were included in downstream analyses.

#### Transcriptome preprocessing

The whole gene RNA read count matrix is used for snmC2T-seq transcriptome analysis. Cells are filtered by the number of genes expressed > 200 and genes are filtered by the number of cells expressed > 10. The count matrix X is then normalized per cell and transformed by ln(X + 1). After log transformation, we use the *scanpy.pp.highly_variable_genes* to select the top 3000 genes based on normalized dispersion, using a process similar to the selection of highly variable methylation features. The selected feature matrix is scaled to unit variance and zero mean per feature followed by PCA calculation.

#### Chromatin accessibility (NOMe-seq) preprocessing

For clustering analysis, cytosine methylation in the GCY context (GmCY) is counted as the open chromatin signal from NOMe-seq, and the HCY context is used to estimate the endogenous mCH background. Highly methylated regions (in GCY context) at a single-cell level are identified by three continuous GmCY sites with maximum distance < 500bp. By applying the same threshold to HCY contexts (non-substrate for GpC methylase), we have determined that the empirical false discovery rate is less than 5%. The binarized peak signal (1 for having highly methylated region in this bin, 0 for no signal or missing data) is then called for all the 5kb nonoverlapping chromosome bins as the NOMe matrix. Similar to snATAC-seq analysis, we then used the SnapATAC (Fang et al., 2019) package to select bins, perform cell-cell Jaccard matrix calculation, and normalize the coverage impact as described in the snATAC clustering analysis section. The PCs are then calculated from the normalized Jaccard matrix.

### General strategies for clustering and manifold learning

#### Consensus clustering on concatenated PCs

We used a consensus clustering approach based on multiple Leiden-clustering (Traag et al., 2018) over K-Nearest Neighbor (KNN) graph to account for the randomness of the Leiden clustering algorithms. After selecting dominant PCs from PCA in all available modalities of different technologies (mCH, mCG for snmC-seq and snmC-seq2; mCH, mCG, RNA, NOMe-seq for snmC2T-seq, etc.), we concatenated the PCs together to construct KNN graph using scanpy.pp.neighbors. Given fixed resolution parameters, we repeated the Leiden clustering 200 times on the KNN graph with different random starts and combined these cluster assignments together as a new feature matrix, where each single Leiden result is a feature. We then used the outlier-aware DBSCAN algorithm from the scikit-learn package to perform consensus clustering over the Leiden feature matrix using the hamming distance. Different epsilon parameters of DBSCAN are traversed to generate consensus cluster versions with the number of clusters that range from minimum to the maximum number of clusters observed in the 200x Leiden runs. Each version contains a few outliers that usually fall into three categories: 1. Cells located between two clusters that have gradient differences instead of clear borders, e.g. L2-3 IT to L4 IT; 2. Cells with a low number of reads that potentially lack information in important features to determine the exact cluster. 3. Cells with a high number of reads that are potential doublets. The number of type 1 and 2 outliers depends on the resolution parameter and is discussed in the choice of the resolution parameter section, the type 3 outliers are very rare after cell filtering. The final consensus cluster version is then determined by the supervised model evaluation.

#### Supervised model evaluation on the clustering assignment

For each consensus clustering version, we performed a Recursive Feature Elimination with Cross-Validation (RFECV) (Guyon et al., 2002) process from the scikit-learn package to evaluate clustering reproducibility. We first removed the outliers from this process, then we held out 10% of the cells as the final testing dataset. For the remaining 90% of the cells, we used tenfold cross-validation to train a multiclass prediction model using the input PCs as features and *sklearn.metrics.balanced_accuracy_score* (Brodersen et al., 2010) as an evaluation score. The multiclass prediction model is based on BalancedRandomForestClassifier from the imblearn package that accounts for imbalanced classification problems (Lemaître et al., 2017). After training, we used the 10% testing dataset to test the model performance using the balanced_accuracy_score score. We kept the best model and corresponding clustering assignments as the final clustering version. Finally, we used this prediction model to predict outliers’ cluster assignments, we rescued the outlier with prediction probability > 0.5, otherwise labeling them as outliers.

#### Choice of resolution parameter

Choosing the resolution parameter of the Leiden algorithm is critical for determining the final number of clusters. We selected the resolution parameter by three criteria: 1. The portion of outliers < 0.05 in the final consensus clustering version. 2. The final prediction model performance > 0.95. 3. The average cell per cluster ≥ 30, which controls the cluster size in order to reach the minimum coverage required for further epigenome analysis such as DMR calling. All three criteria prevent the over-splitting of clusters thus we selected the maximum resolution parameter under meeting the criteria using grid search in each specific clustering analysis below.

#### Cluster marker gene identification and cluster trimming

After clustering, we used a one-vs-rest strategy to calculate methylation (methyl-marker) and RNA (rna-marker, for snmC2T-seq only) marker genes for each cluster. We used all the protein coding and long non-coding RNA genes with evidence level 1 or 2 from gencode v28. For the rna-marker, we used the *scanpy.tl.rank_genes_group* function with the Wilcoxon test and Benjamini-Hochberg multi-test correction, and filtered the resulting marker gene by adjusted P-value < 0.01 and log2(fold-change) > 1, we also used AUROC score as a measure of marker gene’s predictability of corresponding cluster, and filtered genes by AUROC > 0.8. For the methyl-marker, we used the normalized gene body mCH rate matrix to calculate markers for neuronal clusters and the normalized gene body mCG rate matrix for non-neuronal clusters, and we modified the original Wilcoxon test function to used a reverse score to select genes that have significant decrease (hypomethylation). Marker gene is chosen based on adjusted P-value < 0.01, delta methylation level change < −0.3 (hypo-methylation), AUROC > 0.8. The delta methylation level is calculated as the normalized methylation rate change between the cluster and the mean value of the rest clusters.For the ensemble methylome clustering, if cluster with the number of methyl-markers < 10 is detected, the cluster with the minimum total marker genes are merged to the closest clusters based on cluster centroids euclidean distance in the PC space, then the marker identification process is repeated until all clusters found enough marker genes.

#### Manifold learning

The T-SNE and UMAP embedding are run on the PC matrix the same as the clustering input using the scanpy package.

### Identification of open chromatin regions using snmC2T-seq GmCY profiles

The methylation level of each GCY site was normalized to the average GCY cytosine methylation level of the surrounding 100kb region. GCY sites with normalized methylation greater than 2.5 were considered significantly methylated (GmCY) sites. The density of GmCY sites across the genome was modeled using Poisson distribution by MACS2 (Zhang et al., 2008) and regions with significant enrichment of GmCY sites were identified with MACS2 *callpeak* with p-value < 0.01. Peaks with q-value < 0.01 were selected for downstream analyses.

### snATAC-seq data generation

Combinatorial barcoding single nucleus ATAC-seq was performed as described previously in Fang et al. (Fang et al., 2019). Isolated brain nuclei were pelleted with a swinging bucket centrifuge (500 x g, 5 min, 4°C; 5920R, Eppendorf). Nuclei pellets were resuspended in 1 ml nuclei permeabilization buffer (5 % BSA, 0.2 % IGEPAL-CA630, 1mM DTT and cOmplete^TM^, EDTA-free protease inhibitor cocktail (Roche) in PBS) and pelleted again (500 x g, 5 min, 4°C; 5920R, Eppendorf). Nuclei were resuspended in 500 µL high salt tagmentation buffer (36.3 mM Tris-acetate (pH = 7.8), 72.6 mM potassium-acetate, 11 mM Mg-acetate, 17.6% DMF) and counted using a hemocytometer. Concentration was adjusted to 4500 nuclei/9 µl, and 4,500 nuclei were dispensed into each well of a 96-well plate. For tagmentation, 1 μL barcoded Tn5 transposomes (Fang et al., 2019) were added using a BenchSmart™ 96 (Mettler Toledo), mixed five times and incubated for 60 min at 37 °C with shaking (500 rpm). To inhibit the Tn5 reaction, 10 µL of 40 mM EDTA were added to each well with a BenchSmart™ 96 (Mettler Toledo) and the plate was incubated at 37 °C for 15 min with shaking (500 rpm). Next, 20 µL 2 x sort buffer (2 % BSA, 2 mM EDTA in PBS) were added using a BenchSmart™ 96 (Mettler Toledo). All wells were combined into a FACS tube and stained with 3 µM Draq7 (Cell Signaling). Using a SH800 Fluorescence-activated cell sorter (Sony), 40 nuclei were sorted per well into eight 96-well plates (total of 768 wells) containing 10.5 µL EB (25 pmol primer i7, 25 pmol primer i5, 200 ng BSA (Sigma). Preparation of sort plates and all downstream pipetting steps were performed on a Biomek i7 Automated Workstation (Beckman Coulter). After addition of 1 µL 0.2% SDS, samples were incubated at 55 °C for 7 min with shaking (500 rpm). 1 µL 12.5% Triton-X was added to each well to quench the SDS. Next, 12.5 µL NEBNext High-Fidelity 2× PCR Master Mix (NEB) were added and samples were PCR-amplified (72 °C 5 min, 98 °C 30 s, (98 °C 10 s, 63 °C 30 s, 72°C 60 s) × 12 cycles, held at 12 °C). After PCR, all wells were combined. Libraries were purified according to the MinElute PCR Purification Kit manual (Qiagen) using a vacuum manifold (QIAvac 24 plus, Qiagen) and size selection was performed with SPRI Beads (Beckman Coulter, 0.55x and 1.5x). Libraries were purified one more time with SPRI Beads (Beckman Coulter, 1.5x). Libraries were quantified using a Qubit fluorometer (Life technologies) and the nucleosomal pattern was verified using a Tapestation (High Sensitivity D1000, Agilent). The library was sequenced on a HiSeq2500 sequencer (Illumina) using custom sequencing primers, 25% spike-in library and following read lengths: 50 + 43 + 37 + 50 (Read1 + Index1 + Index2 + Read2) (Preissl et al., 2018).

### snATAC-seq data processing

Using a custom python script, we first de-multicomplexed FASTQ files by integrating the cell barcode (concatenate reads pair in I1.fastq and I2.fastq) into the read name (R1.fastq and R2 fastq) in the following format: “@”+“barcode”+“:”+“original_read_name”. Demulticomplexed reads were aligned to the corresponding reference genome (hg19) using bwa (0.7.13-r1126) (Li and Durbin, 2009) in pair-end mode with default parameter settings. Alignments were then sorted based on the read name using samtools (v1.9) (Li et al., 2009). Pair-end reads were converted into fragments and only those that are 1) properly paired (according to SMA flag value); 2) uniquely mapped (MAPQ > 30); 3) with length less than 1000bp were kept. Since fragments were sorted by barcode (integrated into the read name), fragments belonging to the same cell (or barcode) were automatically grouped together which allowed for removing PCR duplicates for each cell separately. Using remaining fragments, a snap-format (Single-Nucleus Accessibility Profiles) file was generated. snap file is hierarchically structured hdf5 file that contains the following sessions: header (HD), cell-by-bin matrix (BM), cell-by-peak matrix (PM), cell-by-gene matrix (GM), barcode (BD) and fragment (FM). HD session contains snap-file version, date, alignment and reference genome information. BD session contains all unique barcodes and corresponding meta data. BM session contains cell-by-bin matrices of different resolutions (or bin sizes). PM session contains cell-by-peak count matrix. PM session contains cell-by-gene count matrix. FM session contains all usable fragments for each cell. Fragments are indexed for fast search. A detailed documentation of snap file can be found here (https://docs.google.com/document/d/1AGyn_WJTr0A1SKcfrEgum-jvAJjWd84jZ6DwbwiGRaQ/edit?usp=sharing). After generating the SNAP file, we filtered cell barcodes based on the following criteria 1) Total Sequencing Fragments (>1,000); 2) Mapping Ratio (>0.8); 3) Properly Paired Ratio (>0.9); 4) Duplicate Ratio (<0.5); 5) Mitochondrial Ratio (<0.1). (Fang et al., 2019).

### Clustering analysis of snATAC-seq data

We used the snapATAC package for the clustering analysis of snATAC-seq data, the detail steps were described in (Fang et al., 2019). Briefly, we used the binarized cell-by-bin matrix of the whole genome 5kb non-overlapping bins as input (1 means open, 0 means close or missing data). We first the coverage of each bin and converted the coverage distribution to log-normal distribution and converted the bin coverage to z-score. Bins with extremely high (zscore > 1.5) or low coverage (zscore < −1.5), or overlap with ENCODE blacklist (Amemiya et al., 2019) are removed. We then converted the cell-by-bin matrix into cell-by-cell similarity matrix by calculating the Jaccard index between cells. To normalize the cell coverage impact on Jaccard index, we used the observed over expected (OVE) method from snapATAC, which calculate the residual of the linear regression model between expected Jaccard matrix given cell coverage and the overserved matrix. We then perform PCA on standardized residual matrix and used top 25 PCs for leiden clustering (resolution = 1) and UMAP visualization.

### Open chromatin peak calling using snATAC-seq data

Open chromatin peaks were identified using snATAC-seq reads combined for each cell type using MACS *callpeak* with the following parameters -f BED --nomodel --shift 37 --ext 73 --pvalue 1e-2. Peaks with q-value < 0.01 were further selected for downstream analyses.

### snRNA-seq data generation

Nuclei were isolated from human postmortem brain tissues and sorted based on NeuN fluorescence as previously described (Hodge et al., 2019). Each sample contained approximately 80% NeuN-positive and 20% NeuN-negative nuclei. snRNA-seq data was generated using 10x Genomics v3 single cell chemistry per the manufacturer’s protocol. RNA-seq reads were aligned with Cell Ranger v3 using the human GRCh38.p2 reference genome, and intronic and exonic mapped reads were included in gene expression quantification.

### snRNA-seq clustering and annotation (Used in Fig 1, 3, 6)

Nuclei were included in downstream analysis if they passed the following QC thresholds: > 500 genes detected (UMI > 0) in non-neuronal nuclei or > 1000 genes detected (UMI > 0) in neuronal nuclei; and doublet score < 0.3. Cells were grouped into transcriptomic cell types using the iterative clustering procedure described in (Tasic et al., 2018). Briefly, genes from the mitochondrial and sex chromosomes were excluded, and expression was normalized to UMI per million and log2-transformed. Nuclei were clustered using the following steps: high variance gene selection, dimensionality reduction, dimension filtering, Jaccard–Louvain or hierarchical (Ward) clustering, and cluster merging. Differential gene expression (DGE) was computed for every pair of clusters, and pairs that did not meet the DGE criteria were merged. Differentially expressed genes were defined using two criteria: 1) significant differential expression (> 2-fold; Benjamini-Hochberg false discovery rate < 0.01) using the R package limma and 2) binary expression (CPM > 1 in more the half of cells in one cluster and < 30% of this proportion in the other cluster). We define the deScore as the sum of the −log10(false discovery rate) of all differentially expressed genes (each gene contributes to no more than 20), and pairs of clusters with deScore < 150 were merged. This process was repeated within each resulting cluster until no more child clusters met DGE or cluster size criteria (minimum of 10 cells). The entire clustering procedure was repeated 100 times using 80% of all cells sampled at random, and the frequency with which nuclei co-cluster was used to generate a final set of clusters, again subject to differential gene expression and cluster size termination criteria. Clusters were identified as outliers if more than 40% of nuclei co-expressed markers of inhibitory (GAD1, GAD2) and excitatory (SLC17A7) neurons or were NeuN+ but did not express the pan-neuronal marker SNAP25. Median values of total UMI counts and gene counts were calculated for each cluster and used to compute the median and inter-quartile range (IQR) of all cluster medians. Clusters were also identified as outliers if the cluster median QC metrics deviated by more than three times the IQRs from the median of all clusters. In total, 23,379 nuclei passed QC criteria and were split into three broad classes of cells (13,997 excitatory neurons, 7,094 inhibitory neurons, and 1,914 non-neuronal cells) based on NeuN staining and cell class marker-gene expression. A final merge step required at least 4 marker genes to be more highly expressed in each pair of clusters. The clustering pipeline is implemented in an R package publicly available at github (https://github.com/AllenInstitute/scrattch.hicat). The clustering method is provided by the run_consensus_clust function.

## FIGURE-SPECIFIC METHODS

### Cell line dataset analysis (Figure S1)

#### Clustering

For both scmCT-seq and snmCT-seq cell line datasets, PCA was used for the dimension reduction of the mCG and RNA matrices. Since only two cell types (H1 and HEK293T) need to be separated, only the first 5 PCs from each matrix were selected to construct K-Nearest Neighbor (KNN) graphs (K=25). On each KNN graph for mCG and RNA, Leiden clustering (r=0.5) is used to determine the two clusters and tSNE was used to visualize the PCs. Clusters were annotated by examining the genome-wide methylation levels and marker gene expression. Data acquired from single cells or nuclei were then merged for each cluster for comparisons with bulk methylome and transcriptome data.

#### Comparison to bulk H1 and HEK293 Methylome

The bulk HEK293 cell whole-genome bisulfite sequencing (WGBS-seq) data were downloaded from Libertini et. al. (GSM1254259) (Libertini et al., 2015). The bulk WGBS-seq data of the H1 cell was downloaded from Schultz et al (GSE16256) (Schultz et al., 2015). Methylpy was used to call CG-DMRs between these two cell lines(Schultz et al., 2015). DMRs were filtered by DMS (differentially methylated sites) ≥ 5 and methylation level difference ≥ 0.6.

#### Bulk H1 and HEK293 RNA Data Analysis

The bulk HEK293 cell RNA-seq data was downloaded from Aktas et. al. (GSE85161) (Aktaş et al., 2017), the bulk H1 cell RNA-seq data was downloaded from encodeproject.org (ENCLB271KFE, generated by Roadmap Epigenome). Gene count tables and bigwig tracks were generated using human GENCODE v19 gene annotation.

### snmC2T-seq baseline clustering (Used in Fig 1)

To perform clustering analysis on the human frontal cortex snmC2T-seq dataset only, we first preprocessed three modalities separately as described in the preprocessing section above. We then concatenate all the dominant PCs together to run the consensus clustering identification (resolution = 1). We annotated the clusters based on marker genes reported in the previous studies (Hodge et al., 2019; Luo et al., 2017). We also calculated the UMAP coordinates based on concatenated PCs (Figure 1D) and PCs from every single modality separately (Figure S2H-J).

#### Methylome ensemble clustering (Used in Fig 2, 3, 4, 5, 6, 7)

To generate an ensemble cell type taxonomy for the human frontal cortex, we combine four methylome based technologies (Figure 2A, snmC2T-seq, snmC-seq, snmC-seq2, sn-m3C-seq) in this study. Due to the high cell-type diversity, we performed a two-level iterative clustering analysis.

##### Level 1 clustering to identify major cell types

We first preprocessed the methylation matrix as described above for each technology separately to obtain the corresponding highly variable feature matrix. We then used scanorama to integrate all cells using the union of highly variable features from all technologies, with K=25 and default values for other parameters. After the integration, we performed PCA on the integrated matrix and used the dominant PCs for the subsequent consensus clustering analysis (resolution = 0.5) as described above. We also calculated UMAP coordinates using the ensemble PCs (Figure 2C).

##### Level 2 clustering to identify subtypes for each major cell type

After level 1 clustering, we selected cells from each major cell type and repeated all the steps from highly variable feature selection to final clustering (K=20, resolution = 0.8) including scanorama integration. The highly variable features selected in this step are more specific to intracluster diversity of each major type, which helps to better separate the subtype. The subcluster UMAP coordinates are calculated from PCs in each subtype analysis (Figure 2C insets).

### snmC2T-seq - snRNA-seq integration (Figure 3, excitatory, inhibitory, non-neuron separately)

To perform the integration analysis of snmC2T-seq transcriptome and snRNA-seq, we separate the cells into three broad classes: excitatory neurons, inhibitory neurons, and non-neuronal cells. The RNA features used for the integration by Scanorama come from two sources for each cell class: 1) highly variable genes across individual cells; 2) cluster level RNA marker genes. To validate that the cluster level RNA marker genes are relevant for neuronal processes, we performed a synapse-specific GO enrichment test using the SynGO terms and all brain expressed genes as background (Koopmans et al., 2019). The −log(adjusted P-value) of SynGO biological process enrichment in each selected gene set is color-coded on the sunburst chart of the hierarchical SynGO terms (Figure 3C,3D).

We then used the union of RNA features found in snmC2T-seq transcriptome and snRNA-seq for Scanorama integration and PCA calculation. The dominant PCs were then used to perform a co-clustering analysis on the cells profiled by snmC2T-seq cell or snRNA-seq. Instead of directly using the co-clustering results, we used this intermediate clustering assignment to calculate the overlap score between the original methylome ensemble clusters and the snRNA-seq clusters. The overlap score range from 0 to 1 is defined as the sum of the minimum proportion of samples in each cluster that overlapped within each co-cluster (Hodge et al., 2019), a higher score between one methylome cluster and one snRNA-seq cluster indicate they consistently co-clustered within one or more co-clusters.

### Integration of DNA methylome and snATAC-Seq data (Figure 2)

Ensemble methylomes and snATAC-seq data from neurons and glia were integrated separately using our recently developed Single Cell Fusion method (see section **Computational data integration with SingleCellFusion**). The top 4000 variable genes across clusters in the snmC2T-seq and snATAC-seq data were identified using a Kruskal-Wallis test; 1,652 genes were identified as being variable in both datasets and were used for the subsequent integration. For snATAC-seq the gene body was extended to include the promoter region (2kb upstream TSS). Prior to integration mCH and open chromatin levels at gene bodies were smoothed to reduce sparseness (k = 20, ka = 4, epsilon = 1, p = 0.9; see section **Within-modality smoothing**) using a diffusion based smoothing method adapted from MAGIC ((van Dijk et al., 2018)). A constrained k-nearest neighbors graph was generated among cells across 2 datasets (k=20, z=10; see section **Cross-modality imputation by Restricted k-Partners**). Instead of calculating Euclidean distance in reduced dimensions, here we simply used Spearman correlation across 1,652 genes as the distance measure between cells. We used the kNN graph to impute the gene body mCH profile for each ATAC-Seq nucleus. The observed (ensemble methylomes) and imputed (snATAC-Seq nuclei) gene body mCH was then jointly used for Leiden clustering and UMAP embedding. Each snATAC-seq nucleus was assigned to a major cell type if at least half of it’s restricted k-Partners belonged to that cell type, remaining cells were removed from subsequent analysis (n=499, 3.98%).

### Cross-validation of cell clusters (Figure 4)

The analysis starts with 2 cell-by-gene data matrices: one for gene-body non-CG DNA methylation (mCH) and the other for RNA expression. We first filter out low-quality cells and low-coverage genes. After removing glia and outliers in the snmC2T-Seq dataset, we get 3,898 high-quality neuronal cells. By selecting genes expressed in >1% of cells and with >20 cytosines coverage at gene body in >95% of cells, we get 13,637 sufficiently covered genes. Then we normalize the mCH matrix by dividing the raw mCH level by the global mean mCH level of each cell; and we normalize the RNA matrix by (log_10_(TPM+1)).

The goal of cluster cross-validation is to cluster cells with one part of the features, and to validate clustering results with the other part of features. We first generate clusterings with different granularity, ranging from coarse to very fine, using DNA methylation features. Clusterings are generated by the Leiden method applied to the top 20 principal components with different settings of the resolution parameter controlling granularity. Following clustering, we randomly split cells into training set and test set. Using the training set, we estimate the cluster centroids of RNA expression. Using the test set, we calculate the mean squared error (MSE) between the RNA expression profile of individual cells and that of cluster centroids. This procedure can be reversed by clustering with RNA features and evaluation with DNA methylation features.

To summarize the results, we plot the curve of number of clusters versus the mean squared error. To ensure robustness, clustering is repeated five times with different random seeds, and with 10 repetitions of 5-fold cross-validation procedure on different random splits of training and test set.

### Quantification of over-splitting and under-splitting of cell clusters (Figure 4)

We define cross-validation based scores to quantify over-splitting (S_over_) and under-splitting (S_under_), respectively. Ideally, each cluster should be globally distinct from other clusters (not over-split) and locally homogeneous, with little systematic difference within the same cluster (not under-split). S_over_ measures global uniqueness, i.e. how distinct one cluster is compared with other clusters. S_under_ measures local homogeneity, i.e. how similar cells within one cluster are to each other. These scores are designed based on cross-validation between transcriptomic and epigenomic information of snmC2T-Seq, as we attempt to address how well clusters defined by one data modality (transcriptome or DNA methylation) can be validated by the complementary data modality.

Over- and under-splitting scores are computed based on a cross-modality distance measure. This distance measure treats the two data modalities as independent measurements, as if they came from separate DNA methylation and transcriptome assays performed on independent groups of cells. Given a cell-by-gene mCH matrix *X* (raw mCH level normalized by global mCH level of each cell), and a cell-by-gene RNA-expression matrix *Y* (log_10_(TPM+1)), we simultaneously reduce the dimensionality of both matrices by canonical correlation analysis (Butler et al., 2018):

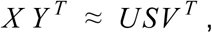

where *U* and *V* are cell-by-feature orthogonal matrices (feature dimension = 20) of the canonical coefficients of cells measured by mCH and RNA, respectively. S is a diagonal matrix of the canonical correlations. We define cross-modality distance between cell *i* (measured by mCH) and *j* (measured by RNA) as the Euclidean distance in the low dimensional feature space:

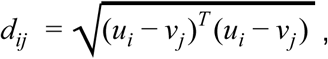

Where *u_i_* and *v_j_* are the *i*’ th column of *U* and the *j*’ th column of *V*, respectively. We then build a bipartite graph, connecting each cell’s profile in one modality with the k-nearest neighbors in the other modality. We refer to the these cross-modality neighbors as “k-partners,” 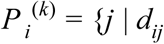 are the k smallest distances for cell *i* }.

Next, we take advantage of the multimodal measurements to define over- and under-clustering metrics using the cross-modality partner cells. The over-clustering score, *S_over_*, is defined as one minus the mean fraction of each cell’s k-partners (with *k* = |*C_i_*| = number of cells in cluster *i*) that are from the same cluster:

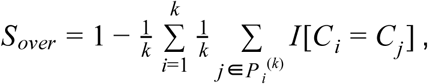

Where *I*[] is the indicator function and *C_i_* is the cluster label of cell *i*. Thus, if all of a cell’s k-partners share the same cluster label, *S_over_* = 0, while larger values of *S_over_* indicate less cross-modality stability for the clusters.

To quantify under-splitting, we compared each cell’s cross-modality distance to itself, *d_ii_*, with the distance to other cells in the same cluster. We define each cell’s self-radius, *r_i_*, as the maximum k value such that all k-partners are closer than *d_ii_*:

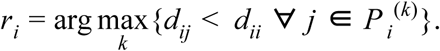

In other words, a cell’s self-radius is the number of *k*-partners whose cross-modality distance is less than the cell’s cross-modality distance to itself.

We compared each self-radius *r* to the size of the cluster. For an ideal cluster with no internal heterogeneity, its cells’ self-radii are uniformly distributed between 0 and the size of the cluster, because all its cells are equivalent to each other. We proved this empirically with simulated clusters generated by randomly shuffling gene features within each cluster (pink line in Figure 4F). For an under-split cluster, its cells’ self-radii are much smaller than the cluster size, indicating it can be potentially further split into sub-clusters. We therefore define *S_under_* as the number of cells whose self-radius was ≤ 25% of the cluster size |*C_i_*|, normalized by |*C_i_*|/4. For an ideal cluster, this score should be 1; for an under-split cluster, it should be greater than 1.

### Computational data integration with SingleCellFusion (Figure 4)

Several computational methods have been proposed for integrating multiple single cell sequencing datasets across batches, sequencing technologies, and modalities(Butler et al., 2018; Haghverdi et al., 2018; Hie et al., 2019; Korsunsky et al., 2019; Welch et al., 2019). Many of these methods share a basic strategy of identifying neighbor cells across datasets. However, existing methods have not been optimized to integrate single cells from multiple transcriptomic and epigenomic data modalities, with potentially large systematic differences in the features measured for each dataset. Here, we integrated the transcriptomes and DNA methylomes of the snmCT-Seq dataset, treating the two data modalities as if they were acquired by two independent single-modality experiments in different cells. We developed a new data integration method, *SingleCellFusion*, for this task (available at: https://github.com/mukamel-lab/SingleCellFusion), which is based on finding k-partners, i.e. nearest neighbors across data modalities (see previous section). Nearest neighbor based data integration has been successfully applied to combine multiple RNA-Seq datasets (Haghverdi et al., 2018; Hie et al., 2019), while other approaches including canonical correlation analysis (CCA) and non-negative matrix factorization (NMF) have previously been used for integrating transcriptomic and epigenomic data (Butler et al., 2018; Welch et al., 2019). Single Cell Fusion is designed to robustly integrate DNA methylation, ATAC-Seq and/or RNA-Seq data. The procedure comprises 4 major steps: preprocessing: within-modality smoothing, cross-modality imputation, and clustering and visualization.

1. **Preprocessing.** We define a gene-by-cell feature matrix for both transcriptomes and epigenomes. Transcriptomic features are log_10_(TPM+1) normalized. DNA methylation data is represented by the mean gene body mCH level, normalized by the global (genome-wide) mean mCH level for each cell. We select genes with significantly correlated gene body mCH and RNA expression (FDR < 0.05) across neuronal cells as features (n=5,107 genes).
2. **Within-modality smoothing.** To reduce the sparsity and noise of feature matrices, we share information among cells with similar profiles using data diffusion (van Dijk et al., 2018). First, we generate a kNN graph of cells based on Euclidean distances in PC space [ndim = 50, k=30]. We next construct a sparse weighted adjacency matrix *A*. We first apply a Gaussian kernel on the distance between cell i and cell j: 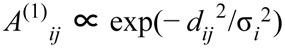, where σ*_i_* is the distance to the *k_a_* -th [*k_a_* =5] nearest neighbor of cell i. We set diagonal elements to zero, *A*^(1)^*_ii_* = 0, and also set all elements to zero if they are not part of the kNN. We then symmetrize the matrix, *A*^(2)^ = *A*^(1)^ + *A*^(1)^*^T^*, and normalize each row: *A*^(3)^*_ij_* = *A*^(2)^/*a_i_*, where 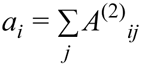. Finally, we reweight the adjacency matrix with a parameter, *p*, that explicitly controls the relative contribution of diagonal and non-diagonal elements: *A* = *p I* + (1 − *p*) *A*^(3)^, where *I* is the identity matrix. We chose p=0.9 for DNA methylation; p=0.7 for RNA. Finally, we smooth the raw feature matrix by matrix multiplication with the adjacency matrix.
3. **Cross-modality imputation by Restricted k-Partners (RKP)**. Each cell has a set of measured features in one data modality (RNA or mC), which we call the “source modality.” The goal of this step of the analysis is to impute the missing features from the other data type, called the “target modality.” For each cell in the source modality, we select a set of k-partners in the target modality and use the average of the k-partners’ features to estimate the missing modality for the original cell. However, care must be taken to avoid hub cells in the target modality which form k-partner relationships with a large fraction of all cells in the source modality. One way to avoid hub cells is by including only mutual nearest neighbors (MNN) (Haghverdi et al., 2018). We developed an alternative approach, restricted k-partners (RKP), that efficiently finds a set of k-partners for every source modality cell, while ensuring that every target-modality cell is connected with a roughly equal number of source modality cells.

As above, we first reduce the dimensionality of both source and target data matrices by canonical correlation analysis, retaining the top 50 canonical components. We then iterate over all cells in the source modality (in random order) k times, connecting each with its most similar partner cell in the target modality. Whenever a target modality cell is partnered with more than *k*′ source modality cells, we remove it from the pool of eligible target cells so that it will not be the partner of additional source cells. We set *k*′ = [*z k N_source_*/*N_target_*]_+_, where and *z* ≥ 1 is a relaxation parameter that determines how much variability in the number of partners is allowed across target modality cells and []_+_ is the ceiling function. If *z* = 1 then every target cell will be connected to exactly *k*′ or *k*′ − 1 cells. We set *z* = 3, meaning that any individual target modality cell can have at most 3 times as many partners as the average. This algorithm is efficient and, in our analyses, provides robust k-partner graphs for cross-modality data imputation.

Having determined each source cell’s restricted k-partners, we next impute the target features by averaging over the smoothed feature vectors of each cell’s k-partners.

4. **Clustering and visualization.** After imputation, we cluster and visualize cells from the 2 data modalities as if they are from the same dataset. We reduce dimensionality for all cells and each set of features (features of each modality) by performing PCA, keeping the top 50 PCs. To balance the contributions of each modality, we divide the features in each modality by their total standard deviation. We concatenate PC matrices as the final feature matrix for downstream embedding and clustering. Next, we perform UMAP embedding (Haghverdi et al., 2018) on the PC matrix [n_neighbors=60, min_dist=0.5]. Finally, we perform Leiden clustering (Traag (Haghverdi et al., 2018) on the kNN graph (symmetrized, unweighted) generated from the final PC matrix [Euclidean distance, k=30, resolution=0.8, 1, 2, 4].

### LIGER integration (Figure S4)

The LIGER integration was performed largely per the recommended RNA-to-methylation integration pipeline. A LIGER object was created for the transcriptome and methylome data from each snmC2T-seq profiled cell, and 5,145 RNA-mCH coupled genes (see **Correlation analysis of RNA expression and gene body DNA methylation**). Transcriptome data was normalized per LIGER’s default function, and mCH was scaled to the maximum. For the non-negative matrix factorization 20 factors (k) were selected with a penalization of 5 (lambda). The data was quantile normalized and a UMAP embedding was generated on the factors.

### Correlation analysis of RNA expression and gene body DNA methylation (Figure 5)

For each gene, we compute the Spearman correlation coefficient between RNA expression (log_10_(TPM+1)) and gene body mCH (raw mCH level normalized by global mCH of each cell) over all neurons or over a subset of cells. To know if a correlation is statistically significant, we randomly shuffled cell labels to generate an empirical null distribution. Significantly correlated genes are defined with empirical FDR < 0.05. Applying this method to 3,898 neurons in snmC2T-Seq dataset, we get 5,145 genes with significant negative correlation between RNA and mCH (RNA-mCH coupled).

### Eta Squared of Genes Across Clusters (Figure 5)

For each gene used for correlation analysis we compute the η^2^ across neuronal sub clusters (n=52) generated from ensemble methylomes (Fig. 2) for both RNA (log_10_(TPM+1)) and gene body mCH (normalized by global mCH of each cell) signals. We also compute η^2^ across 10X RNA-seq clusters for the same genes.

### H3K27me3 ChIP-Seq data processing (Figure 5)

We downloaded published H3K27me3 ChIP-Seq data of purified excitatory and inhibitory neurons from human prefrontal cortex (Kozlenkov et al., 2018). We calculated the average ChIP-Seq signal intensity (RPKM) across the gene body for excitatory and inhibitory neurons.

### Cell type dendrogram and sub-cluster merge along the lineage (Figure 6)

The cell-type hierarchy of inhibitory and excitatory cells were calculated separately using the concatenated PCs from mCG and mCH as the features used for computing cluster centroids. We used *scipy.cluster.hierarchy.linkage* function to calculate the ward linkage. Based on the linkage results, we merged the CpG sites from single-cell ALLC files in 2 steps: 1) we merged the single-cell ALLC files into each of the sub-clusters, 2) we then merge the sub-clusters into all nodes appeared in the dendrograms. The merged CpG ALLC files are then used in the lineage-DMR analysis.

### Neural lineage specific DMR calling and motif enrichment analysis (Figure 6)

We used the *methylpy findDMR* function (Schultz et al., 2015) to identify mCG lineage-DMRs for each pair of lineages using merged ALLC files. The DMRs identified by methylpy in each branch comparison is further filtered by mCG rate difference > 0.3 and the number of differentially methylated sites (DMS) >= 2. Lineage pairs with >10^4^ DMRs identified were used for motif enrichment analysis and TF marker identification. For each of these DMR sets, we use AME (McLeay and Bailey, 2010) to perform motif enrichment (fisher’s exact test) analysis with the motifs’ Position Weight Matrix (PWM) from the JASPAR database (JASPAR2018 CORE Vertebrates) (Khan et al., 2018). The DMRs are length standardized into ±250bp of region center before motif scanning. Tissue-specific DMRs (without brain tissue, and standardized in the same way) from the Roadmap Epigenomics project (Roadmap Epigenomics Consortium et al., 2015; Schultz et al., 2015) were used as the background.

### TF binding preference to methylated motifs (Figure 6)

To further investigate the methylation level impact on the potential TF binding sites, we selected all the mCG DMSs ±25bp regions from the branch-DMRs and ran motif enrichment using motifs identified from the methyl-SELEX experiment (Yin et al., 2017). In each branch pair, we used the left-DMSs as the background of right-DMS to find the right-branch-specific motif and vise Versa. The significant enriched “TF motif - branch” combinations were then intersected with the corresponding branch pair’s DEG and DMG list to infer their gene mCH or RNA specificity.

### Chromatin accessibility (NOMe-seq) analysis of TF binding motifs (Figure 6H-M)

Genome-wide sites matching TF binding motifs (motif matches) identified by methyl-SELEX (Yin et al., 2017) was identified using FIMO 4.11.4 (Grant et al., 2011) with the following parameters --max-stored-scores 500000 --max-strand --thresh 1e-5. Methyl-SELEX only quantified the effect of CpG methylation on TF binding. Therefore only genomics sites containing CG dinucleotides were selected for further analyses. As discussed above, we found the local density of methylated GmCY sites (normalized GmCY level > 2.5 compared to the surrounding 100kb region) as a reliable marker of chromatin accessibility. For each major cell type, the density of GmCY sites was quantified for motif matches that overlap with hypomethylated or hypermethylated DMRs. Figure 6H shows the average chromatin accessibility at motif matches across major cell types. TF binding motifs were ranked by the difference of chromatin accessibility between motif matches located in hypomethylated and hypermethylated DMRs. To test the enrichment of MethylPlus and MethylMinus TFs, the ranked motif list was divided into 5 bins and the enrichment or depletion in each bin was tested using Matlab *hygecdf* function.

### Partitioned heritability analysis (Figure 7)

Bulk human fetal frontal cortex methylomes from a PCW 20 donor (Lister et al., 2013) and a PCW 19 donor (Luo et al., 2016) were previously published. Fetal frontal cortex DMRs were identified using *methylpy findDMR* function (Schultz et al., 2015) by comparing to adult bulk neuronal (NeuN+) and non-neuronal (NeuN-) methylomes (Lister et al., 2013). Fetal brain Dnase-seq samples included fetal day 85d (GSM595922, GSM595923), 96d (GSM595926, GSM595928), 101d (GSM878650), 104d (GSM878651), 105d (GSM1027328), 109d (GSM878652), 112d (GSM665804), 117d (GSM595920) and 142d (GSM665819). Mapped reads files (BED format) were downloaded followed by DNase-seq peak calling using MACS2 2.0.10 with q-value < 0.01. Fetal brain DNase-seq peaks were defined as the union DNase-seq peaks of fetal brain DNase-seq datasets and were supported by at least two samples.

Summary statistics were downloaded from the Psychiatric Genomics Consortium portal (https://www.med.unc.edu/pgc/) for neuropsychiatric trait GWAS - ADHD (Martin et al., 2018), Aggression (Pappa et al., 2016), Anorexia nervosa (Watson et al., 2019), Anxiety (Otowa et al., 2016), ASD (Grove et al., 2019), Bipolar (Stahl et al., 2019), Cognitive Performance (Rietveld et al., 2013), Educational Attainment (Rietveld et al., 2013), Alzheimer’s (Lambert et al., 2013), Internalizing, Loneliness (Gao et al., 2017), Major Depression (Wray et al., 2018), Neuroticism (Smith et al., 2016), OCD (Mattheisen et al., 2015), Schizophrenia (PGC2) (Schizophrenia Working Group of the Psychiatric Genomics Consortium, 2014) and Schizophrenia (PGC1) (Ripke et al., 2013).

The partitioned heritability analysis was performed using LD Score Regression (LDSC) Partitioned Heritability (Finucane et al., 2015). The partitioned heritability analysis was performed by constructing joint linear models by providing multiple regulatory element annotations in addition to the “baseline” annotation. We built a “baseline” annotation using tissue-specific DMRs from non-brain human tissues (Schultz et al., 2015) to control for generic gene regulation characteristics. The reported q-values were derived from the “*Coefficient_z.score*” values reported by LDSC Partitioned Heritability.

## SUPPLEMENTARY FIGURE LEGENDS

**Figure. S1.**
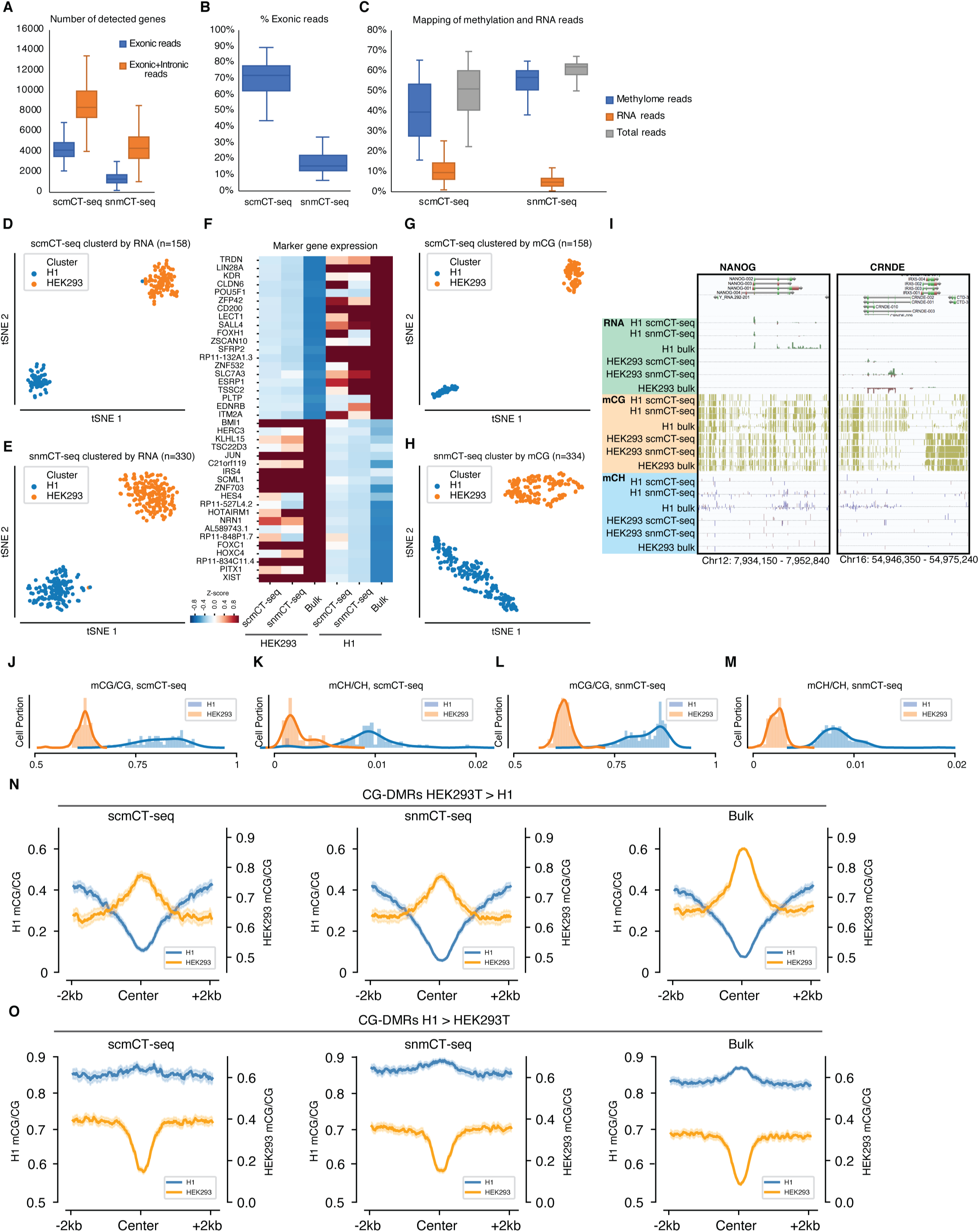
scmCT-seq and snmCT-seq capture transcriptome and DNA methylation signatures of H1 & HEK293 cells. (A-C) Number of detected genes (A), percentage of mapped reads that located in exons (B) and mapping rates of methylation and RNA reads (C) for scmCT-seq and snmCT-seq. (D-E) Separation of H1 and HEK293T cells by tSNE using transcriptome reads extracted from scmCT-seq (D) or snmCT-seq (E) datasets. (F) scmCT-seq and snmCT-seq detect genes specifically expressed in H1 or HEK293T cells. (G-H) Separation of H1 and HEK293T cells by tSNE using DNA methylation information extracted from scmCT-seq (G) or snmCT-seq (H) datasets. (I) Browser view of NANOG and CRNDE loci. (J-M) Distribution of mCG and mCH levels for single H1 and HEK293 cells/nuclei as determined by scmCT-seq and snmCT-seq. (N-O) scmCT-seq and snmCT-seq recapitulate bulk mCG patterns at CG-DMRs showing greater mCG levels in HEK293T (N) or H1 (O) cells.

**Figure S2.**
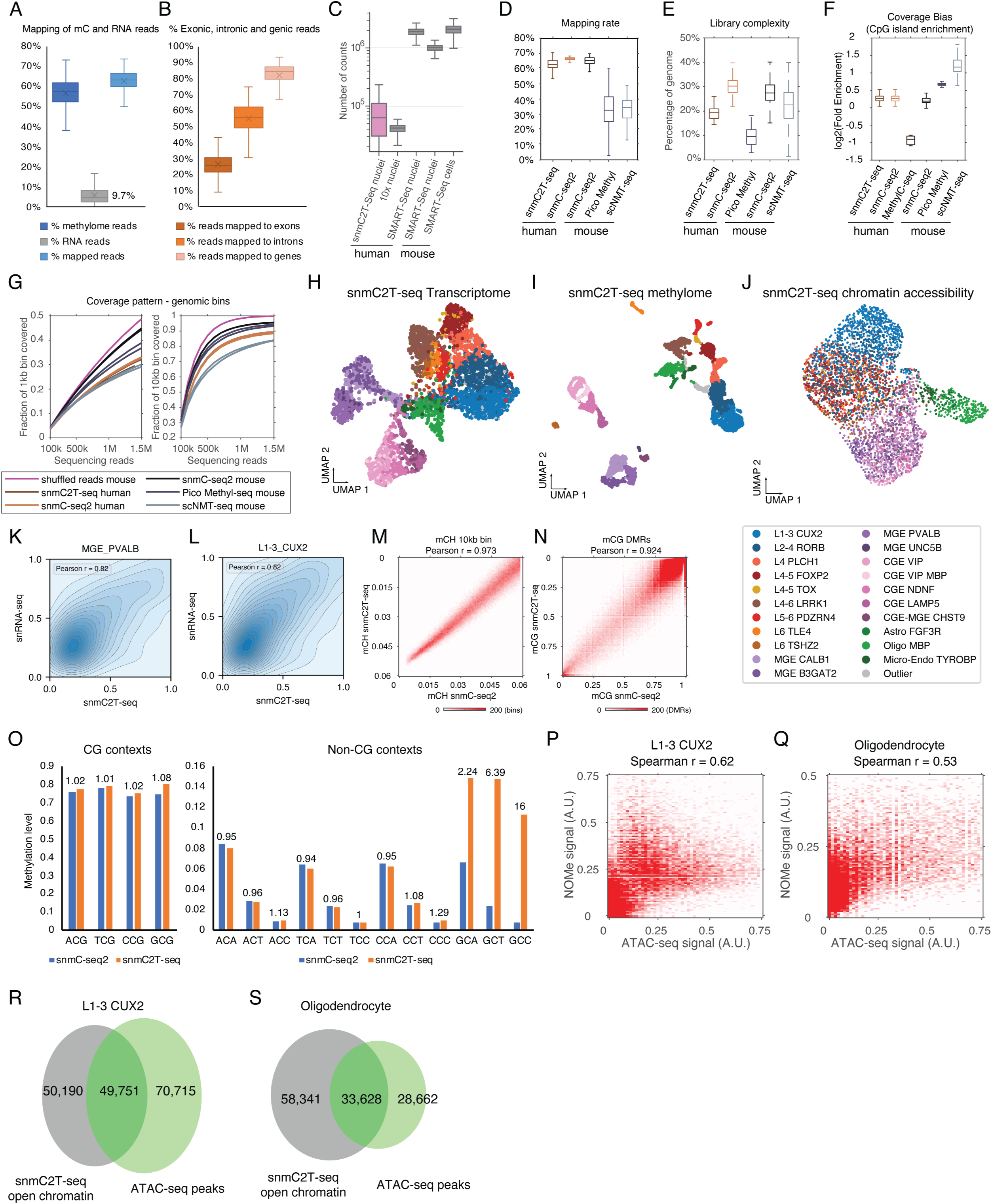
snmC2T-seq generates single-nucleus multi-omic profiles from human brain tissues. (A) The fraction of total snmC2T-seq reads derived from methylome and transcriptome. (B) The fraction of snmC2T-seq transcriptome reads mapped to exons, introns or gene bodies. (C) Boxplot comparing the number of reads detected in each cell/nucleus by different single-cell or single-nucleus RNA-seq technologies. (D-G) snmC2T-seq methylome was compared to other single-cell methylome methods with respect to mapping rate (D), library complexity (E), enrichment of CpG islands (F) and coverage uniformity (G). (H-J) UMAP embedding of 4253 snmC2T-seq cells using single modality information: transcriptome (H), methylome (mCH and mCG, I) and chromatin accessibility (J). (K-L) Pearson correlation of gene expression levels quantified by snmC2T-seq transcriptome and snRNA-seq in MGE PVALB (K) and L1-3 CUX2 (L) cells. (M) Pearson correlation of gene body non-CG methylation quantified with snmC2T-seq methylome and snmC-seq for MGE PVALB cells. (N) Pearson correlation of CG methylation at DMRs quantified with snmC2T-seq methylome and snmC-seq for MGE PVALB cells. (O) Genome-wide methylation level for all tri-nucleotide context (−1 to +2 position) surrounding cytosines shows the sequence specificity of GpC methyltransferase M.CviPI. (P-Q) Spearman correlation between the frequency of methylated GCY sites and ATAC-seq signal at open chromatin sites in L1-3 CUX2 (P) and Oligodendrocyte (Q) cells. (R-S) Overlap of open chromatin peaks identified by snmC2T-seq and snATAC-seq in L1-3 CUX2 (R) and oligodendrocyte (S) cells.

**Figure S3.**
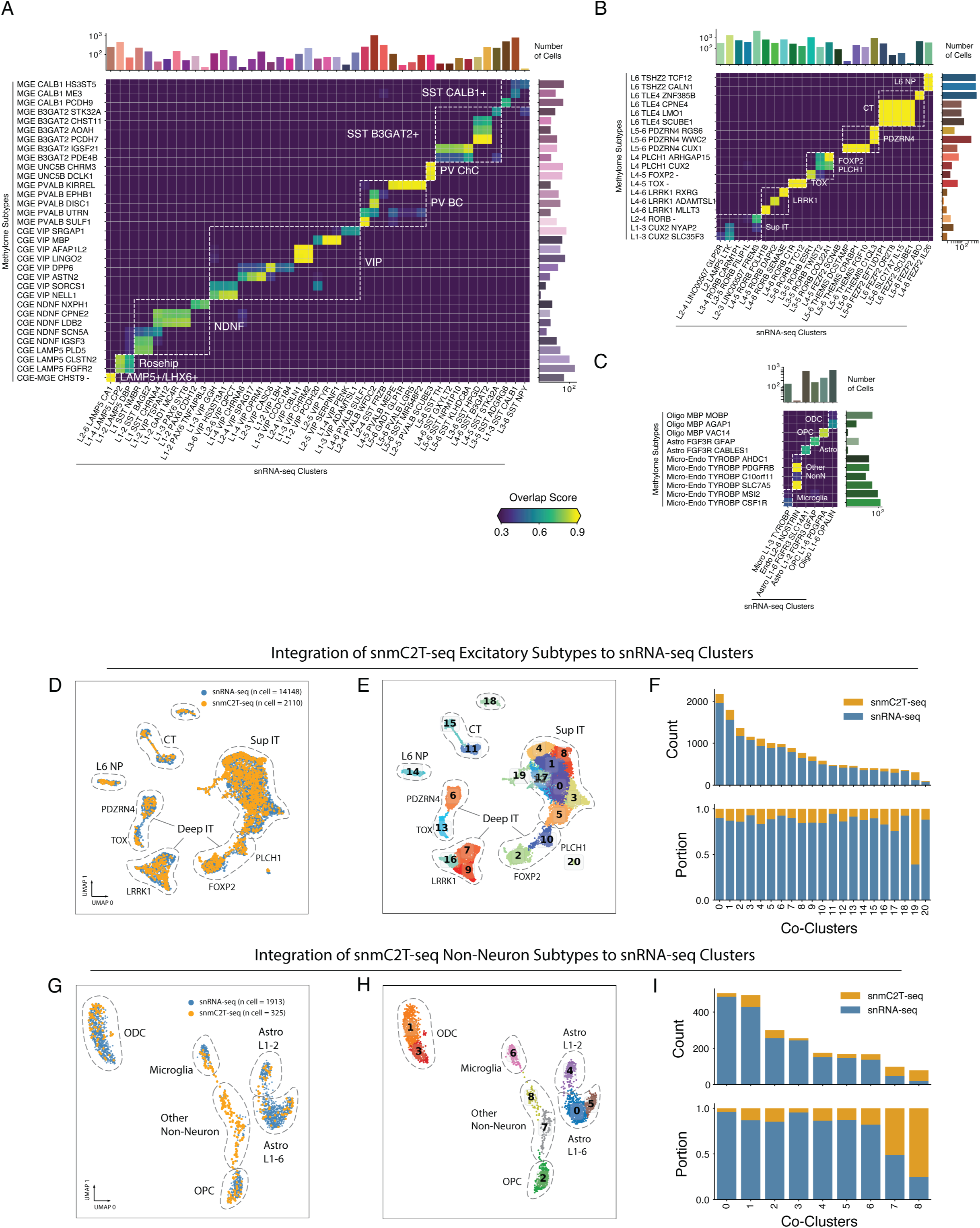
snmC2T-seq recapitulates transcriptome and methylome signatures of neuronal subtypes. (A) Confusion matrix showing the overlap scores between inhibitory subtypes identified by ensemble methylome analysis and snRNA-seq. Known inhibitory cell type groups are annotated by boxes. The upper bar plot indicates the snRNA-seq cell counts per cluster; the right bar plot indicates the snmC2T-seq cell counts per cluster. (B-C) Confusion matrix showing the overlap scores between methylome ensemble subtypes and the original snRNA-seq inhibitory clusters for excitatory neuron clusters (B) and non-neuron clusters (C). Known cell type groups are annotated by boxes. The upper bar plot annotates the snRNA-seq cell counts per cluster, the right bar plot annotates the snmC2T-seq cell counts per cluster. (D-E) UMAP embedding of all excitatory neurons profiled by snmC2T-seq and snRNA-seq after MNN-based integration, colored by technology (D) and joint clusters (E). Known cluster groups are also circled and annotated on UMAP. (F) The composition of cells profiled by snmC2T-seq and snATAC-seq in excitatory neurons joint clusters. The upper and lower bar plots show the counts and portion of cells profiled by the two technologies in each joint cluster, respectively. (G-H) UMAP embedding of all non-neuronal cells from snmC2T-seq and snRNA-seq after integration, colored by technology (G) and joint clusters (H). (I) The composition of cells profiled by snmC2T-seq and snATAC-seq in non-neuronal cell joint clusters.

**Figure S4.**
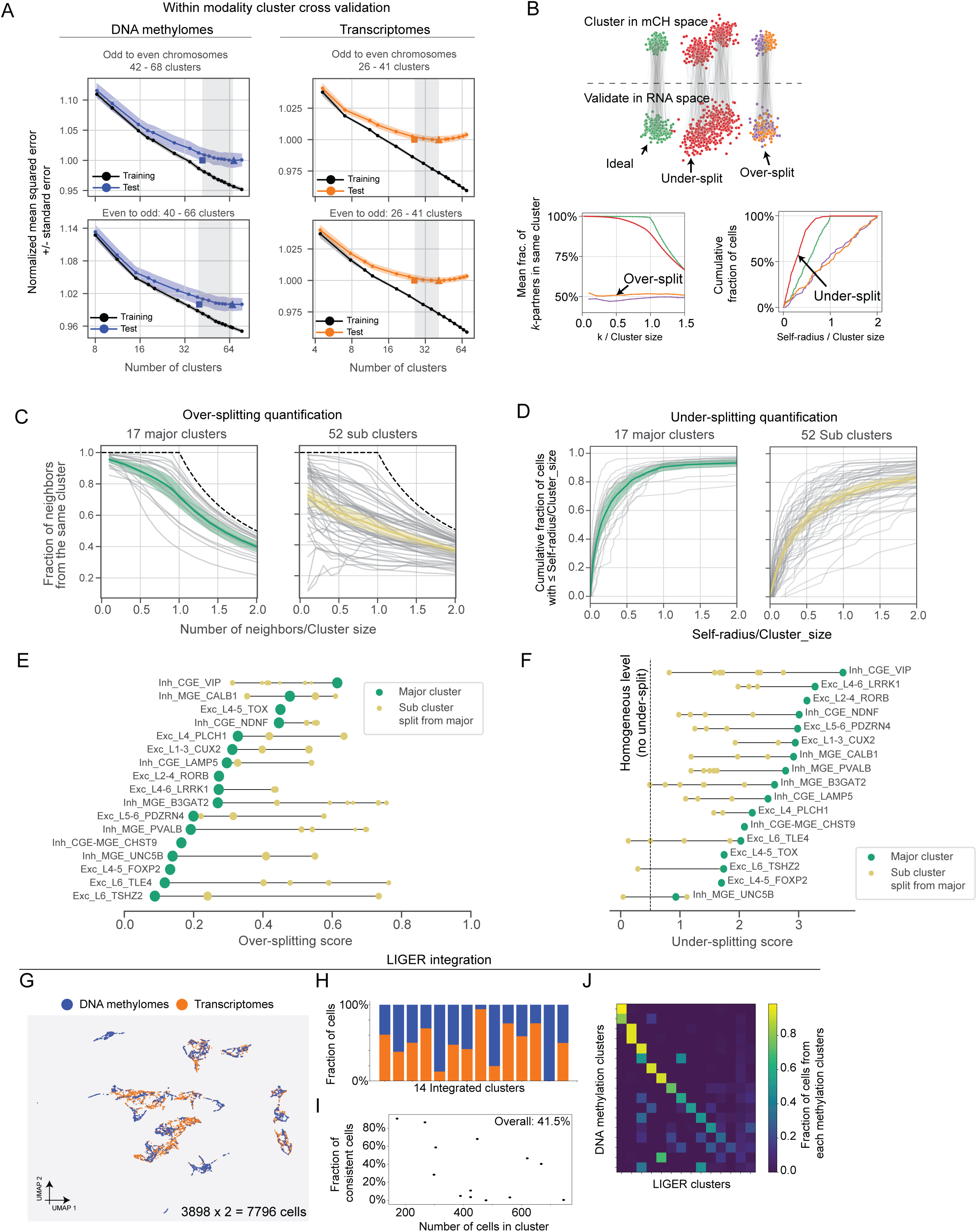
Evaluation of cluster quality with paired transcriptome and methylome profiles. (A) Intra-modality cross-validation of mC- or RNA-defined clusters. Line plots show mean squared error (MSE) between single-cell profile and cluster centroid as a function of the number of clusters. Black lines indicate training error; purple or orange lines indicate test error. Points corresponding to minimum MSE and minimum MSE + 1 standard error are marked by arrows. For the analysis using snmC2T-seq mC information (left panels), gene body mCH profiles of odd (even) chromosomes were used for clustering whereas even (odd) chromosomes were used for testing. Similar analysis was performed using snmC2T-seq transcriptome information (right panels). (B) Schematic diagram of the over- and under-splitting analysis using matched single-cell methylome and transcriptome profiles, complementing Figure 4D. (C) Over-splitting quantification of mC-defined major clusters (n=17) and subclusters (n=52) were quantified by the fraction of cross-modal neighbors found in the same cluster defined by RNA. (D) Under-splitting of clusters was quantified as the cumulative distribution function of normalized self-radius for mC-define major clusters and subclusters. For (C-D), gray lines represent individual clusters while colored lines represents means and confidence intervals. (E) Over-splitting score for each major cluster (in green) and associated sub-clusters (in yellow). Dot size of sub clusters represents cluster size normalized by the size of their “mother” major cluster. (F) Under-splitting score for each major cluster (in green) and associated sub-clusters (in yellow). (G) Joint UMAP embedding of snmC2T-seq transcriptome and mC profiles using the LIGER method, treating the two data modalities as generated from independent datasets. (H) Barplot showing the fraction of cells contributed by each data modality for each integrated cluster. (I) Scatter plot showing the fraction of consistent cells and the size of joint clusters. (J) Confusion matrix of LIGER cluster versus DNA methylation clusters. Values are normalized by each row.

**Figure S5.**
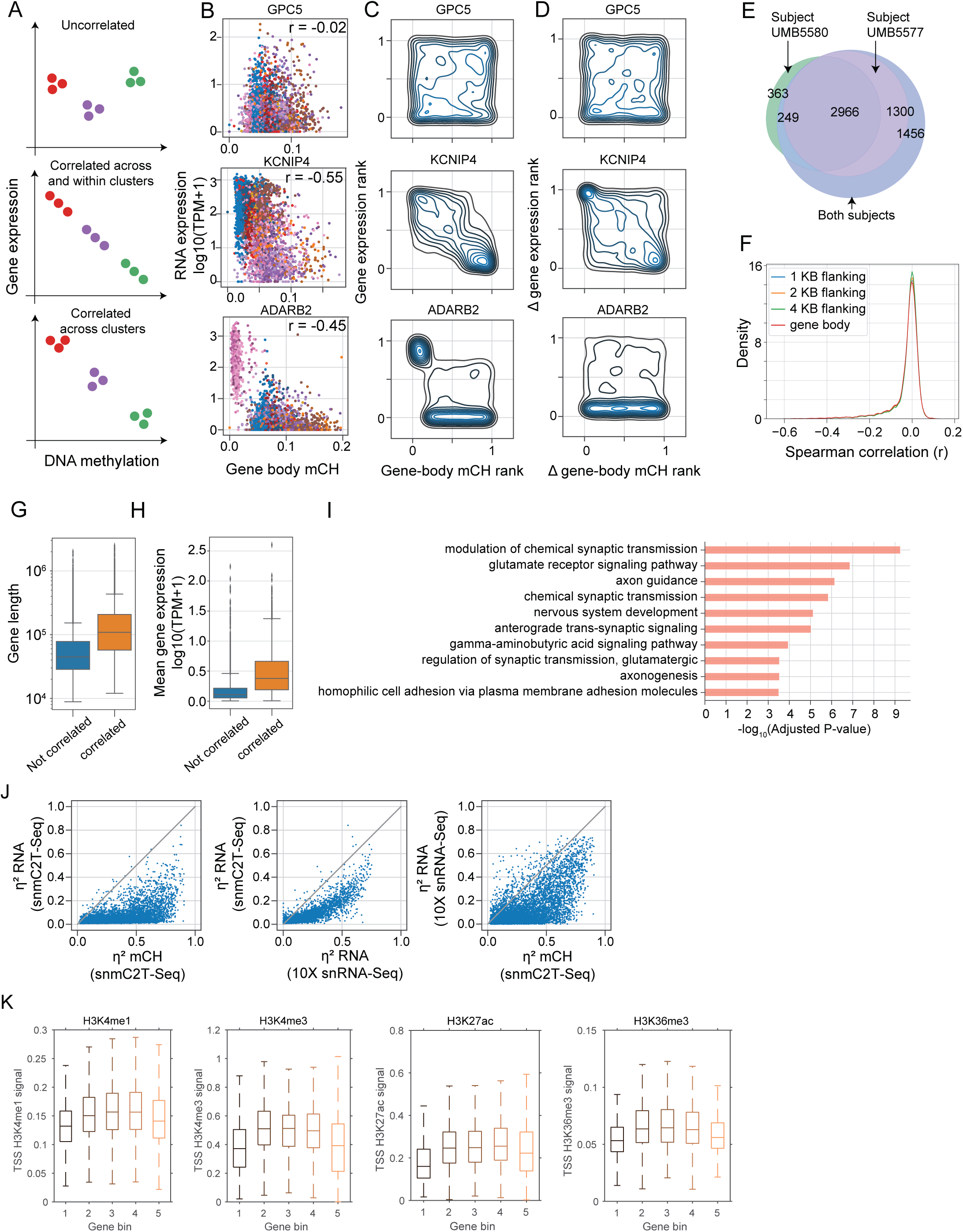
Diverse correlations between gene expression and gene body mCH. (A) Schematic diagram of 3 types of genes with different correlation between gene expression and DNA methylation. (B) Scatter plots of gene body mCH (unnormalized) and gene expression of example genes (KCNIP4, ADARB2, GPC5) across all neuronal cells. Cells are colored by major cell types defined in Figure 2. (C) Contour density plot of gene body mCH rank versus gene expression rank for 3 example genes: GPC5, KCNIP4, and ADARB2. (D) Contour density plot of delta gene body mCH rank (rank of gene body mCH - cluster mean gene body mCH) versus delta gene expression rank (rank of gene expression - cluster mean gene expression) for 3 example genes: GPC5, KCNIP4, and ADARB2. (E) Venn diagram of 3 gene sets--genes significantly correlated between gene body mCH and gene expression called using cells from subject UMB5580, subject UMB5577, and from both subjects. (F) Distribution of spearman correlation coefficient between gene expression and gene level mCH quantified at gene body (red), gene body + 1 kilo-base upstream (blue), + 2 kilo-base upstream (orange) and + 4 kilo-base upstream (green). (G) Boxplot of gene length versus uncorrelated and correlated genes. (H) Boxplot of mean gene expression log10(TPM+1) versus uncorrelated and correlated genes. (I) Gene ontology enrichment of correlated genes. (J) Scatter plots comparing the fraction of variance explained by cell type (η^2^) for each gene from different datasets or data modalities: RNA (from snmC2T-Seq), mCH (from snmC2T-Seq) and 10X (snRNA-Seq from 10X protocols). (K) Boxplots of the distribution of different histone marks at TSS over 5 gene bins grouped according to gene expression ratio of early fetal (PCW 8-9) to adult (>2 yrs).

**Figure S6.**
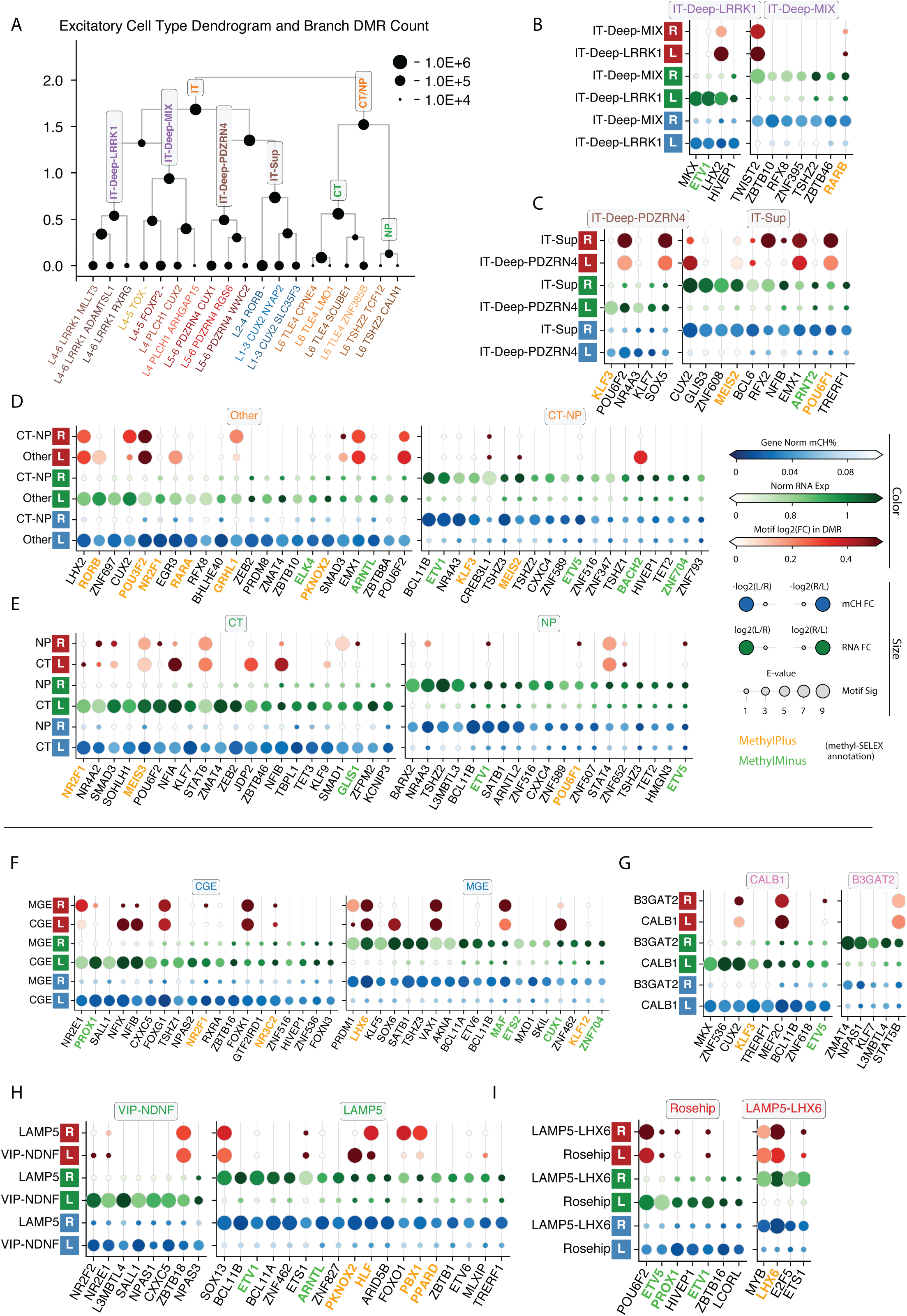
TF binding motif enrichment across human cortical neuronal hierarchy. (A) Excitatory neuron subtype dendrogram. The node size represents the number of DMRs detected between the left and right branches. (B-E) Dot plots view for TFs showing lineage specific motif enrichment, expression and gene body mCH between lineages: IT-Deep-LRRK1 vs IT-Deep-MIX (B); IT-Deep-PDZRN4 vs IT-Sup (C); IT vs CT-NP (D); CT vs NP (E). Colors for every two rows from bottom to top: lineage mean gene body mCH level, lineage mean expression log(1 + CPM), TF motif enrichment log2(fold change). Sizes for every two rows from bottom to top: relative fold change of mCH level from this branch to the other, relative fold change of expression level, E-value of the motif enrichment test. Colors for the motif names: TF motif methylation preference annotated by methyl-SELEX experiment (Yin et al., 2017), orange indicate MethylPlus, green indicate MethylMinus. (F-I) Dot plot view for TFs showing lineage specific motif enrichment, expression and gene body mCH between CGE vs. MGE (F), CALB1 vs. B3GAT2 (G), VIP/NDNF vs. LAMP5 (H) and Rosehip vs LAMP5-LHX6 (D).

**Figure S7.**
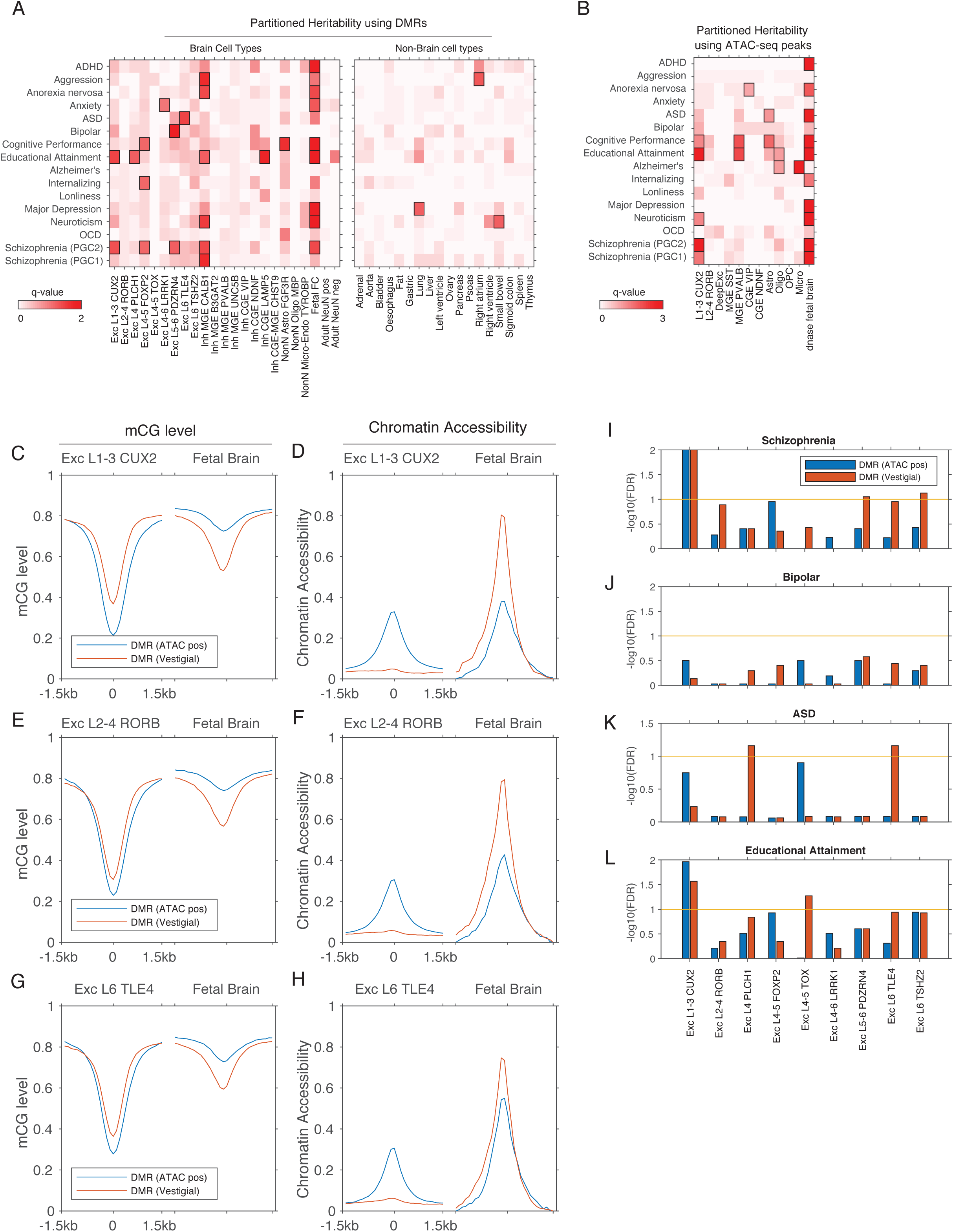
Prediction of causal cell types for neuropsychiatric traits using partitioned heritability analysis. (A) Partitioned heritability analysis using adult brain cell-type specific DMRs and bulk tissue DMRs for fetal cortex, adult NeuN+ population, adult NeuN-population and non-brain tissues. (B) Partitioned heritability analysis using adult brain cell-type specific ATAC-seq peaks and fetal brain DNase-seq peaks. (C-H) CG methylation and chromatin accessibility profiles at adult brain regulatory elements [DMR (ATAC-pos)] and vestigial enhancers. (I-L) Partitioned heritability analysis comparing adult regulatory elements and [DMR (ATAC-pos)] and vestigial enhancers.

## SUPPLEMENTARY TABLES

**Table S1.** Metadata for scmCT-seq data generated from H1 and HEK293 cells

**Table S2.** Metadata for snmCT-seq data generated from H1 and HEK293 cells

**Table S3.** Metadata for human brain specimens

**Table S4.** Metadata for snmC2T-seq data generated from human brain frontal cortex

**Table S5.** Cluster labels of the snRNA-seq dataset

**Table S6.** Cluster labels of single-cell methylomes

**Table S7.** Cluster labels of the snATAC-seq dataset

**Table S8.** Marker genes of brain cell types identified using gene body mCH

**Table S9.** Differentially methylated regions (DMRs) of major brain cell types

**Table S10.** ATAC-seq peaks of major brain cell types

